# SurA is a “Groove-y” Chaperone That Expands Unfolded Outer Membrane Proteins

**DOI:** 10.1101/2019.12.17.878660

**Authors:** Dagan C. Marx, Ashlee M. Plummer, Anneliese M. Faustino, Taylor Devlin, Michaela A. Roskopf, Mathis J. Leblanc, Henry J. Lessen, Barbara T. Amann, Patrick J. Fleming, Susan Krueger, Stephen D. Fried, Karen G. Fleming

## Abstract

The periplasmic chaperone network ensures the biogenesis of bacterial outer membrane proteins (OMPs) and has recently been identified as a promising target for antibiotics. SurA is the most important member of this network both due to its genetic interaction with the β-barrel assembly machinery complex as well as its ability to prevent unfolded OMP (uOMP) aggregation. Using only binding energy, the mechanism by which SurA carries out these two functions is not well understood. Here we use a combination of photo-crosslinking, mass spectrometry, solution scattering, and molecular modeling techniques to elucidate the key structural features that define how SurA solubilizes uOMPs. Our experimental data support a model in which SurA binds uOMPs in a groove formed between the core and P1 domains. This binding event results in a drastic expansion of the rest of the uOMP, which has many biological implications. Using these experimental data as restraints, we adopted an integrative modeling approach to create a sparse ensemble of models of a SurA•uOMP complex. We validated key structural features of the SurA•uOMP ensemble using independent scattering and chemical crosslinking data. Our data suggest that SurA utilizes three distinct binding modes to interact with uOMPs and that more than one SurA can bind a uOMP at a time. This work demonstrates that SurA operates in a distinct fashion compared to other chaperones in the OMP biogenesis network.

**Significance Statement:** Outer membrane proteins play critical roles in bacterial physiology and increasingly are exploited as antibiotic targets. SurA is the most important chaperone in the OMP biogenesis network and is thought to initiate their folding through an interaction with the BAM complex. We observe an unprecedented expansion of unfolded outer membrane proteins when bound to SurA. This expansion suggests a potential mechanism by which SurA can deliver uOMPs to the BAM complex. In addition, this study highlights the use of an integrative/hybrid structural biology approach and emerging methods to map highly heterogeneous structural ensembles such as that of an unfolded protein bound to a chaperone.

## Introduction

Proteins must fold into their native three-dimensional structures to perform their functions. For some proteins this folding process is spontaneous and requires no exogenous factors; however, many proteins – particularly, membrane proteins – are predisposed to populate misfolded states or aggregates that are not functional and can be toxic to the cell (1–3). To suppress these pathways, chaperone proteins promote efficient protein folding through interactions with nascent, unfolded proteins (termed clients) (4–8).

One chaperone network of particular importance is the outer membrane protein (OMP) biogenesis network in gram-negative bacteria. Following translocation across the inner membrane, this network solubilizes hydrophobic, unfolded OMPs (uOMPs) in the aqueous periplasm and delivers them to the β-barrel assembly machine (BAM) complex, which catalyzes uOMP folding into the outer membrane (9–13). OMPs play several critical roles in bacterial physiology such as nutrient uptake, lipid remodeling, and efflux (14). Recently, the OMP biogenesis pathway has been exploited as a target for the development of antibiotics against gram-negative bacteria because drugs that compromise essential OMP maturation need only cross the fairly porous outer membrane and not the tightly regulated inner membrane (15–17).

The uOMP biogenesis chaperone network is comprised of three proteins: SurA, which has been shown to be the most important protein in the pathway, as well as Skp and FkpA (18–23). SurA handles the majority of the flux of unfolded outer membrane proteins (uOMPs) through the periplasm, and accordingly a Δ*surA* null-strain induces a pronounced envelope stress response (9, 18, 24–29). Eight OMPs of varying size (8–26 transmembrane β-strands) and sequence composition have been identified as SurA clients because their expression is notably decreased in the Δ*surA* strain (14, 28, 30–32).

The mechanism by which SurA binds and solubilizes client uOMPs is currently unknown. The lack of ATP in the periplasm implies that the driving forces for SurA’s function must derive from interactions with its clients. Unlike the other members of the OMP biogenesis network that oligomerize to form cages around uOMPs, SurA lacks an obvious cavity to shield uOMPs from the aqueous periplasm and does not readily oligomerize in solution (9). Instead, SurA has a modular structure with three distinct domains connected by flexible linkers shown in Figure 1A: a primarily alpha helical “core” domain comprised of portions from the N- and C-terminal regions, and two peptidyl prolyl isomerase (PPIase) domains (P1 and P2) (33–36). The orientations of the P1 and P2 domains relative to the core domain have been shown to be dynamic, though how the SurA conformational ensemble contributes to uOMP binding is unclear (22, 37–39). The core domain of SurA is thought to be responsible for the majority of the binding energy to small, eight-stranded uOMPs and alone can complement the Δ*surA* strain of *E. coli* (22, 30, 40–48).

**Figure 1.**
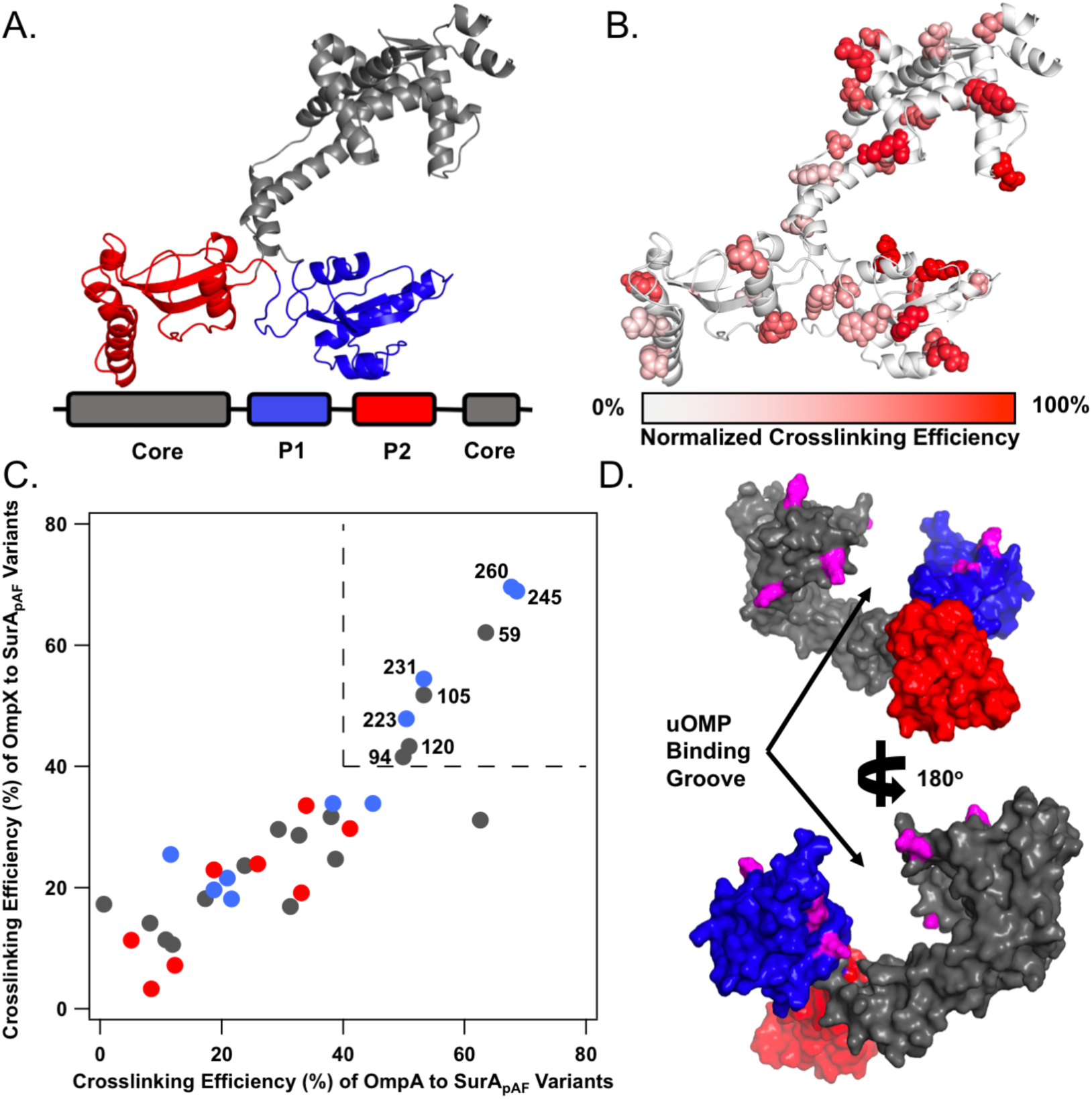
Open SurA binds client uOMPs in a groove between domains. **(A)** The structure of open SurA shown as a cartoon, with the domains colored as depicted in the sequence diagram below (core: gray, P1: blue, P2: red). In this conformation, the three domains of SurA are structurally isolated from each other and do not form extensive inter-domain contacts. **(B)** The 32 surface exposed sites on SurA in which *p*AF was substituted, shown in a space-filling representation. Photo-crosslinking was induced with 5-min UV exposure. Each crosslinking site is colored based on the normalized crosslinking efficiency to uOmpA_171_ as based on quantitative SDS-PAGE (see SI Appendix, Figure S2). The highest crosslinking sites are found on the core and P1 domains of SurA, while P2 exhibits only modest crosslinking efficiency. **(C)** The raw crosslinking efficiencies of SurA to uOmpA_171_ and uOmpX are shown and colored by the SurA domain in which they residue (as in **(A)**). Eight SurA_*p*AF_ variants stand out by having high (>40%) crosslinking efficiency to both uOMP clients, and are labeled with their residue number in the upper right quadrant of the graph (demarcated by dotted lines). **(D)** The eight high efficiency crosslinking sites, shown in magenta, are mapped on to a surface representation of the structure of open SurA. Together, these sites line a groove formed between the core and P1 domains, indicating that uOMPs are primarily bound there.

Structural elucidation of a SurA•uOMP conformation is challenging. As discussed above, SurA has been shown to exist in multiple conformations. Moreover, the unfolded nature of client OMPs poses several additional hurdles that have impeded structural studies using classical techniques: uOMPs lack regular secondary structure, are highly dynamic, and are prone to aggregation (49–52). We address these challenges by capitalizing on the power of an integrative/hybrid structural biology approach that combines data from crosslinking, mass spectrometry, and neutron scattering to elucidate structural features of the SurA•uOMP ensemble. By incorporating photo-crosslinking unnatural amino acids throughout SurA, we find SurA sites with the highest crosslinking efficiencies to clients line a groove formed between the core and P1 domains. Contrast-matching small angle neutron scattering (SANS) of a crosslinked complex reveals that a canonical uOMP client is greatly expanded when bound to SurA. Mass spectrometry analysis of the photo-crosslinked SurA-uOMP complexes identifies specific segments on client uOMPs that preferentially interact with the SurA groove.

Using our experimental data as restraints, we created a sparse ensemble of 40 configurations of SurA•uOMP complexes. We validated structural features present in this ensemble with additional chemical crosslinking mass spectrometry and SANS experiments that were not included in model generation. We identified three distinct uOMP binding modes and higher order stoichiometries that were sufficient to explain the data. Overall, our findings provide a structural basis for how SurA solubilizes its uOMP clients and provide a template for how future studies might elucidate the structures of highly dynamic chaperone complexes with unfolded proteins.

## Results

### SurA crosslinks preferentially with client uOMPs

We identified regions of SurA involved in binding uOMPs using short-distance crosslinkers across the surface of SurA. To accomplish this, we incorporated the unnatural amino acid, *para*-azido-Phenylalanine (*p*AF), at 32 non-conserved, surface-exposed positions on SurA (Figure 1B; see SI Appendix, Fig. S1) using amber suppression (53). *p*AF is a ‘zero-length’ crosslinker, because it forms highly reactive intermediates that crosslink rapidly and non-specifically to any residue within 3–4 Å (though one crosslinking mechanism has been shown to slightly favor reactions with aromatic amino acids) (54, 55). Previously applied in biochemical assays, we report here to the best of our knowledge the first application of *p*AF crosslinking for structural studies (56, 57).

Each SurA variant with a single amino acid substituted for *p*AF (denoted SurA_*p*AF_) was mixed with one of three uOMPs: clients uOmpA_171_ (the transmembrane domain of uOmpA, see SI Methods), uOmpX, or a non-client uOmpLA as a negative control (31, 58). Samples were exposed to UV light for 5 minutes and crosslinking efficiency was measured by quantitative SDS-PAGE (see SI Appendix, Figure S2, Table S1) (59). Crosslinking efficiencies to the two client uOMPs vary dramatically with position on SurA, with the highest efficiency sites all residing on the core and P1 domains (Figures 1B,C). The finding that high-efficiency crosslinking sites for client uOMPs map to the SurA P1 and core domains is consistent with previous experiments that found that removal of the P2 domain did not affect the binding affinity of uOmpA_171_ to SurA (22). In contrast, the non-client uOmpLA showed low crosslinking efficiencies; indeed, only half of the SurA_*p*AF_ variants could form crosslinks with uOmpLA at all (see SI Appendix, Figure S3, Table S1). In addition to the high-efficiency crosslinking sites on cognate client uOMPs, we observed lower levels of crosslinking to client uOMPs at other *p*AF sites across the surface of SurA. The observed differences in crosslinking efficiencies between client and non-client uOMPs indicate that SurA is inherently able to distinguish client uOMP sequences in solution without the aid of other chaperones.

### SurA binds uOMPs in a groove between the core and P1 domains

We sought to identify the uOMP binding site on SurA by visualizing the high-efficiency *p*AF crosslinking sites on known conformations of SurA (see SI Appendix, Table S2)(37, 39). This analysis revealed that residues colocalized around a groove that forms between the core and P1 domains in the open conformation (Figure 1D, see SI Appendix, Fig. S4 and S5). In this conformation, both of the PPIase domains are structurally isolated from the core domain (shown in Figure 1A,B,D).

The uOMP-binding groove is large enough (∼25 × 25 x 25 Å) to shield from water the entire length of either a transmembrane β-strand or β-hairpin of an uOMP (see SI Appendix, Figure S5A). The walls of the groove are formed by the core and P1 domains, which provide hydrophobic patches surrounded by weakly positively charged regions. The base of the groove contains a long hydrophobic stretch (30 Å) and is more positively charged than the walls. Interestingly, the regions of the core and P1 domains outside of the groove are highly negatively charged. The separation of charges on SurA could allow electrostatics to play a role in driving uOMP binding to the groove (see SI Appendix, Figure S5B). In sum, the SurA groove possesses a hybrid chemical nature and size well suited to accommodate the alternating hydrophobic-hydrophilic patterning of uOMP transmembrane domains.

### SurA solubilizes uOmpA_171_ in an expanded conformation

uOMPs are expected to exist in a relatively collapsed, molten globule state in the absence of chaperones (50). This collapsed conformation is maintained when uOmpA_171_ is bound by the other major chaperone in the uOMP biogenesis pathway, Skp (60, 61). To determine whether the overall size and shape of a uOMP changes when bound to SurA, we measured the hydrodynamic properties using small angle neutron scattering (SANS). SANS reports directly on the radius of gyration (*R*_G_) and the maximum end-to-end distance (*D*_Max_) of macromolecules. Moreover, the sample and buffer conditions can be manipulated in a SANS experiment to visualize a selected component within a complex.

We capitalized on this selective contrast feature of SANS and collected the scattering profile of a photo-crosslinked complex composed of protonated SurA_105,*p*AF_ and perdeuterated uOmpA_171_ in 30% D_2_O (Figure 2A). In this experiment, SurA contributes a minor fraction to the scattering contrast (see SI Appendix, Figure S6), and the scattering intensity is primarily contributed by uOmpA_171_. Guinier and P(r) analyses of data collected in this condition revealed that the complex has an *R*_G_ value of 45 ± 3 Å and a *D*_Max_ of 150 ± 10 Å (Figures 2B and C, see SI Appendix, Tables S3 and 4). These sizes far exceed the expected *R*_G_ and *D*_Max_ calculated from the structure of apo-SurA (*R*_G_ = 35 Å; *D*_Max_ = 105 Å). Previous experiments also show that apo-SurA is not denatured by D_2_O (xx). We therefore conclude that the large complex size observed arises from an expanded state of uOmpA_171_ whilst in complex SurA.

**Figure 2.**
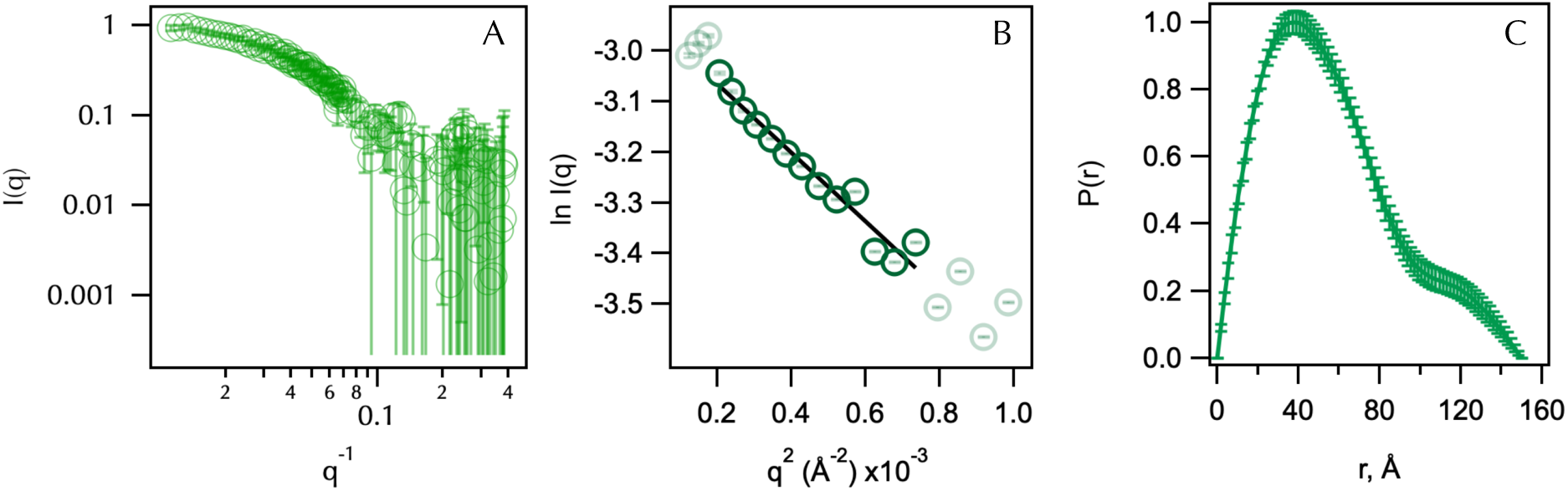
SANS of a SurA_105,*p*AF_-uOmpA_171_ complex reveals an expanded uOmpA_171_. (**A**) Raw scattering profile of protonated-SurA_105,*p*AF_ photo-crosslinked to deuterated-uOmpA_171_ in 30% D_2_O buffer is shown in green. Error bars represent the standard error of the mean with respect to the number of pixels used in the data averaging. (**B**) Linear fit of the Guinier region of the SANS profile determines the *R*_G_ of the complex to be 45 ± 3 Å. (**C**) *P*(*R*) distribution function; D_Max_ is estimated to be 150 ± 10 Å.

To understand the extent to which SurA expands uOmpA_171_ relative to the inherent, unbound uOmpA_171_ molten-globule state, we estimated the intrinsic *R*_G_ and *D*_Max_ of unfolded OmpA_171_. It is impossible to directly measure these parameters with scattering experiments because uOMPs aggregate at the required protein concentrations. However a complementary hydrodynamic parameter of uOmpA_171_, the sedimentation coefficient, *s*, has been previously reported (*s* = 1.65 Svedbergs) (62). We connected *R*_G_, *D*_Max_, and *s*-value using HullRad analysis of atomic models of uOmpA_171_ (63). A series of structural models of unfolded OmpA_171_ were created, and the average *R*_G_ and *D*_Max_ of uOmpA_171_ models that agree with the experimental *s* - value were 25 ± 1 Å and 82.5 ± 9 Å respectively (see SI Appendix, Figures S7 and S8; error is estimated from standard deviations). Thus, the *R*_G_ and *D*_Max_ of uOmpA_171_ are both approximately doubled when it is in complex with SurA. In sum, our *p*AF crosslinking and SANS data support a model where client uOMPs are expanded by SurA: a portion of the client uOMP is bound within the SurA groove, and the remainder of the uOMP is poised to sample transient interactions broadly across the SurA surface.

### SurA preferentially interacts with specific segments on client uOMPs

The expansion of uOMPs by SurA raises the question of where and how they interact with the groove. To further define the molecular basis of the SurA•uOMP interaction, we used photo-crosslinking mass spectrometry (pXL-MS) to identify the segments on uOMPs that crosslinked to the high efficiency SurA_*p*AF_ variants. We identified multiple SurA-binding segments on each client uOMP according to the following criteria.

The eight high-efficiency SurA_*p*AF_ variants were crosslinked to uOmpA_171_ and uOmpX, and subjected to proteolysis with either trypsin only or trypsin and GluC in serial (see Supplementary Appendix). The resulting peptides were analyzed by LC-MS/MS, and crosslinked peptides were identified (with FDR < 0.01) using the MeroX v2.0 software package (see Supplementary Appendix) (64). Summary data of all pXL-MS experiments are given in Supplementary Appendix Table S5 and summary data of all peptide spectrum matches (PSMs) from these pXL-MS experiments are provided as Supplementary Data 1 and 2.

Figure 3A shows the crosslinking pattern identified for both uOmpA_171_ and uOmpX for each high efficiency SurA_*p*AF_ variant. We hypothesized that uOMP residues that repeatedly crosslink to SurA_*p*AF_ variants (>50%) delineate preferred uOMP binding segments. This strategy was necessary because the high reactivity of the nitrene group formed upon photolysis of *p*AF combined with the difficulty of determining the relative abundance of various crosslinked peptides made it impossible to use the crosslink sites from any individual SurA_*p*AF_ variant to distinguish preferred binding segments.

**Figure 3.**
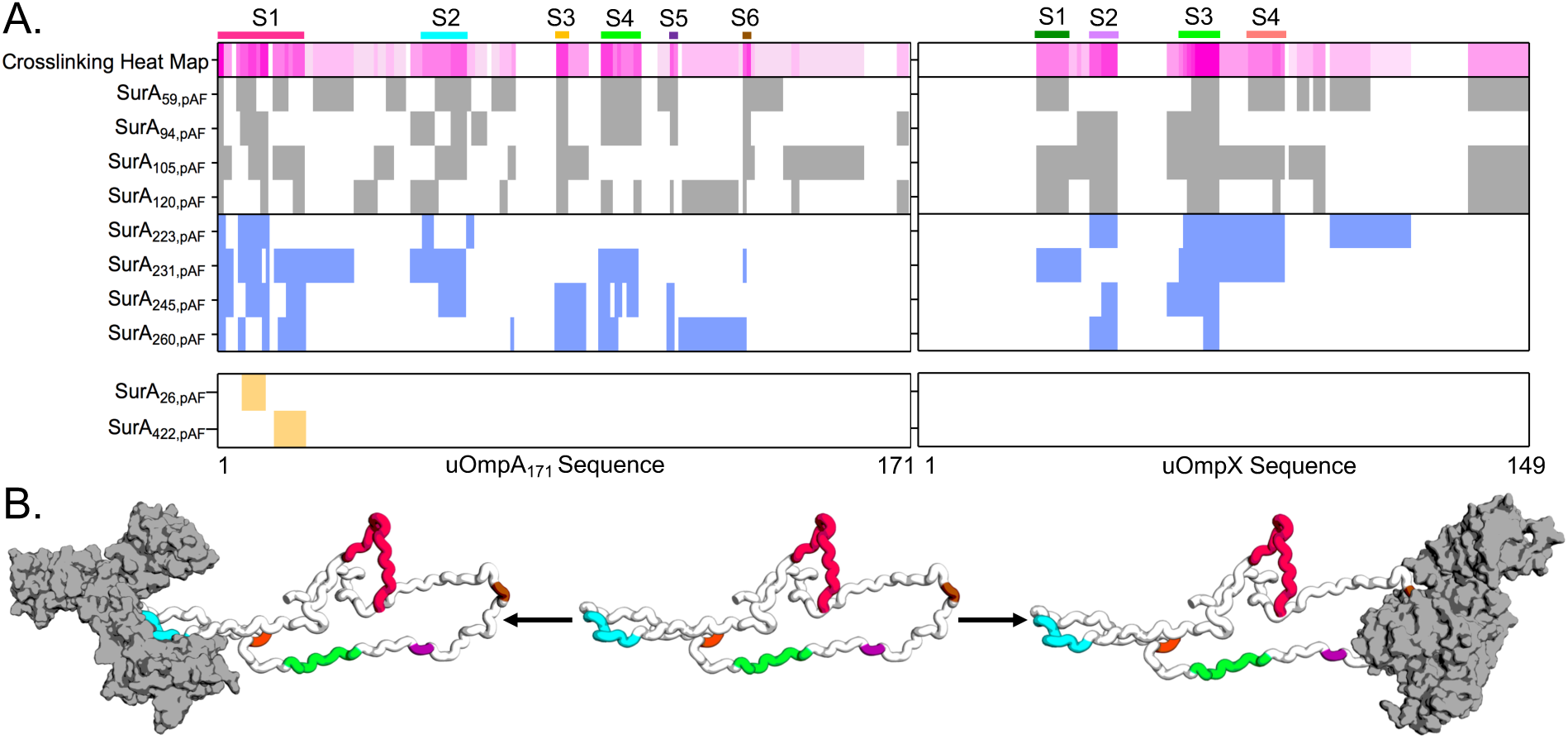
Photo-crosslinking mass spectrometry (pXL-MS) identifies segments on client uOMPs that bind SurA. **(A)** The crosslinking patterns for eight high-efficiency SurA_pAF_ variants are shown to two uOMP clients (uOmpA_171_ and uOmpX). Constructs that place *p*AF on the core domain are colored gray (top register) and constructs that place *p*AF on the P1 domain are colored blue (second register). The crosslinking heat map depicts the frequency a given residue on a client uOMP crosslinks to *p*AF in eight separate crosslinking experiments (darker magenta indicates residue is crosslinked more often). Binding segments are demarcated with a colored bar above the heat map and a label (S1-6 or S1-4). The bottom register shows results from two negative control studies using SurA_26,*p*AF_ and SurA_422,*p*AF_ (see main text for explanation). Only one uOmpA_171_ crosslinked peptide was found for each of these constructs, while uOmpX did not crosslink at all. **(B)** An expanded uOmpA_171_ model with hydrodynamic properties matching the contrast-matched SANS experiment is shown as a cartoon with the SurA-binding segments colored as in **(A)**. Two different segments (S2, left, o1s022 and S6, right, o1s021) are shown bound to SurA (shown in gray with a surface representation), suggesting that more than one copy of SurA could bind a single copy of uOmpA_171_ with minimal steric clash.

Using this criterion, we identified six SurA-binding segments on uOmpA_171_ (residues 1-21, 51-61, 84-86, 95-104, 112-113, and 130-131) and four SurA-binding segments on uOmpX (residues 29-36, 42-48, 64-73, and 81-89) as shown in Figure 3A. The identified uOMP binding segments vary in length and location between the client uOMPs and crosslink to residues on both the core and P1 domains of SurA. The sequences of these SurA-binding segments are unusually enriched in tyrosine residues (10 of the 13 tyrosines on uOmpA_171_ appear in segments; P=0.003 by Chi-square test). Indeed SurA has been shown to preferentially bind to peptides with high aromatic content, affirming our criteria for defining uOMP segments (30). Serendipitously, in OmpA many of these tyrosines are highly conserved according to the Pfam database (Pfam ID: 01389), perhaps indicating importance for these residues in OmpA biogenesis (65).

To validate the importance of the SurA groove as the uOMP binding site, we performed two controls. In the first, we carried out pXL-MS on SurA_422,*p*AF_, which places the *p*AF away from the groove on the opposite side of the core domain. SurA_422,*p*AF_ only crosslinks to a single site on uOmpA_171_ and did not crosslink to uOmpX at all, demonstrating a preference of uOMP interactions with the SurA groove (Figure 3A). Secondly, we found a similarly small number of crosslinks upon mutation of a highly conserved residue in the construct SurA_26,*p*AF_ that also resides away from the groove. Taken together, these experiments show that binding segments on uOMPs selectively distinguish and interact with the SurA groove.

Figure 3B shows a structural model of uOmpA_171_ in an expanded state with a *D*_Max_ equal to the experimentally determined size of uOmpA_171_ when bound to SurA. Each of the putative binding segments are highlighted with the same colors used in Figure 3A. Two possible SurA**•**uOmpA_171_ complexes are shown with SurA bound to the second (left) and last two segments (right) of uOmpA_171_. The presence of multiple SurA-binding segments on uOmpA_171_, along with its expanded size, could allow for more than one copy of SurA to simultaneously bind to different segments of a single uOmpA_171_ (Figure 3B), as further supported by the presence of higher molecular weight bands in SDS-PAGE of some crosslinked SurA_*p*AF_ variants (see SI Appendix, Figure S2).

### Modeling the Structural Features of SurA•uOmpA_171_

Our SDS-PAGE (Figure 1B & SI Appendix Figure S2), SANS (Figure 2), and pXL-MS (Figure 3) experiments support a mechanism in which SurA binds a defined client uOMP segment in the SurA groove. The remainder of the uOMP is greatly expanded and presumed to be dynamic. To visualize this binding mode in more detail, we built 23 models of uOmpA_171_ bound to SurA using our experimental findings as restraints.

Figures S9 and S10 (see SI Appendix) provide flowcharts describing the process by which these models were generated and where each piece of experimental information was included; computational modeling is described further in SI Appendix. In essence, we docked the binding segments of uOmpA_171_ identified by pXL-MS to the groove of SurA using distance restraints generated by HADDOCK (66). The uOmpA_171_ component of these models were then expanded to be compatible with the SANS data (see SI Appendix, Figures S11A and B).

The Hill coefficient reported for SurA binding uOmpA_171_ is greater than 1, indicating more than one copy of SurA can interact with uOmpA_171_ at a time (23). This multiplicity is also consistent with our finding of higher molecular weight bands in SurA_*p*AF_ SDS-PAGE experiments. Accordingly, we also created 17 models with additional SurA protomers docked to the expanded uOmpA_171_ (see SI Appendix, Figure S11 C-E). Hydrodynamic parameters of each of the 40 structural models created are tabulated in Table S6.

### Chemical crosslinking validates features of SurA•uOmpA_171_ models, and reveals distinct binding modes

To validate structural features of the SurA•uOmpA_171_ sparse ensemble, we performed XL-MS with the chemical crosslinker disuccinimidyl dibutyric urea (DSBU) on WT SurA and our client uOMPs. In total, we identified 46 unique DSBU crosslinks between SurA and uOmpA_171_ and 17 unique DSBU crosslinks between SurA and uOmpX (see SI Appendix, Figure S12, Table S5, and Supplementary Data 3–5).

To ascertain underlying similarities in our structural models, we performed spectral biclustering on a matrix of all solvent-accessible surface distances (SASDs, calculated using JWalk) for the 46 identified crosslinks in all 23 structural models of SurA•uOmpA_171_ (see SI Appendix, Figure S13 shows this analysis, Supplementary Data 6 provides all SASDs and their associated scores) (67–69). Due to the conformational heterogeneity and multiplicity of the SurA•uOmpA_171_ sparse ensemble, no single structural model captured all of the DSBU crosslinks. However, three distinct SurA•uOmpA_171_ binding modes emerged from this analysis, which are each explained by a unique subset of the identified crosslinks.

Figure 4A shows an example of the first binding mode, which is found in seven of our SurA•uOmpA_171_ models, wherein segment 1 is bound in the groove and the remainder of uOmpA_171_ is projecting away from SurA. The second binding mode, shown in Figure 4B, is present in five SurA•uOmpA_171_ models and is similar to binding mode one, except segment 2 is bound in the groove. Seven members of the sparse ensemble evinced a more complex topology in which the uOMP threads through the groove several times, defining a third binding mode (Figure 4C). Similar to the first binding mode, segment 1 is bound in the groove, but in this latter mode segments 3, 4, and 5 now make extensive contacts with the groove and P1 on a “second pass.” Together, these three binding modes of the SurA•uOmpA_171_ complex cover 75% (34 out of 46; see SI Appendix, Figure S13) of the identified DSBU crosslinks and help explain why so many regions of uOmpA_171_ can crosslink with the SurA groove (Figure 3).

**Figure 4.**
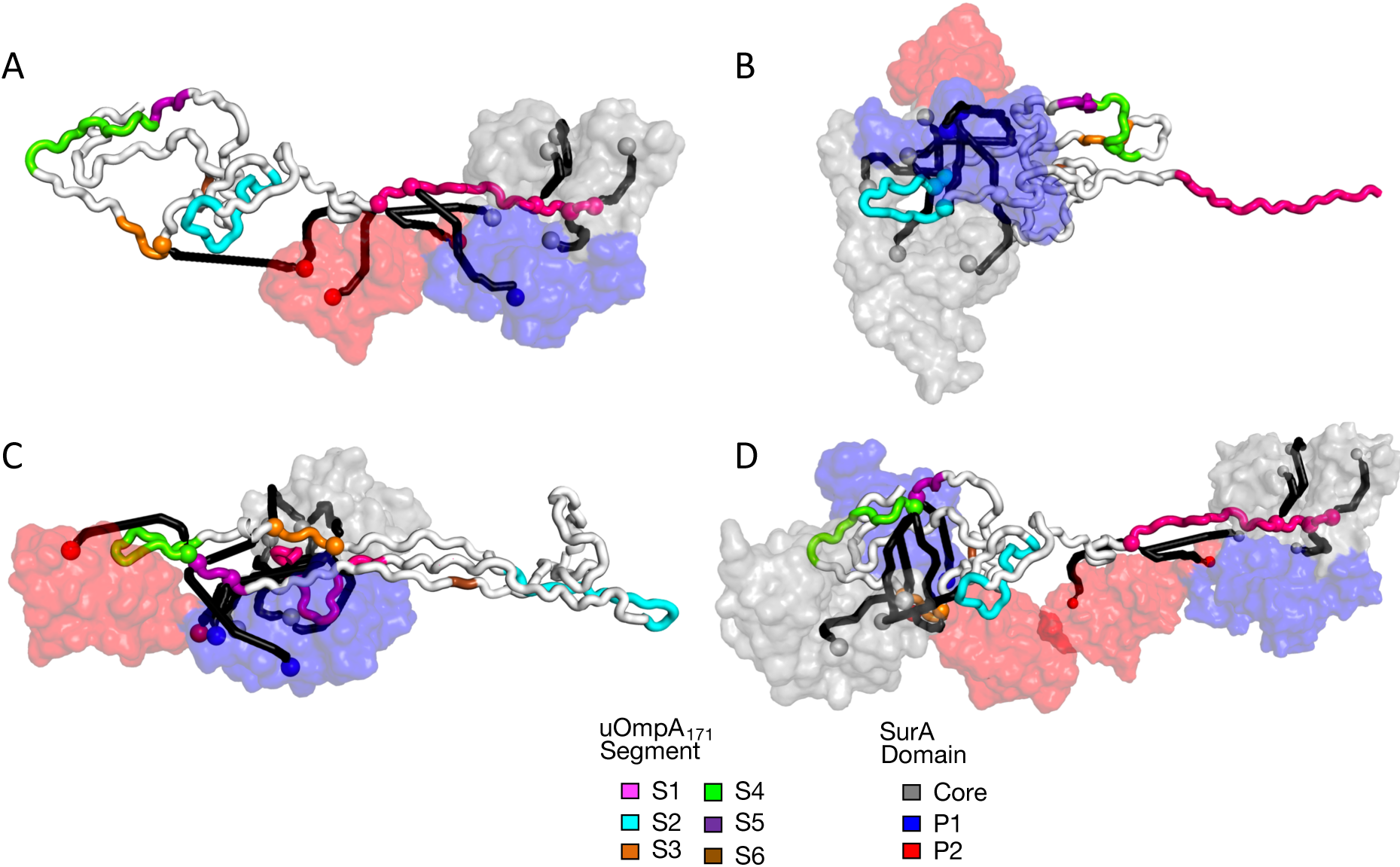
Structural models of three urA•uOmpA_171_ binding modes validated by XL-MS and SANS. SurA is represented as a surface (colored as in Figure 1A) and uOmpA_171_ is represented as a tube (with binding segments demarcated by color as in Figure 3). DSBU crosslink sites are represented as spheres with crosslinks depicted as SASDs created by Jwalk. Crosslinks captured in each model are colored black (see SI Appendix, Figure S13 for details). **(A)** A representative structural model of the first binding mode (o1s016), supported by a cluster of ten crosslinks primarily between S1 (pink) and the core and P2 domains of SurA. **(B)** A representative structural model of the second binding mode (o1s010), supported by a cluster of nine crosslinks primarily between S2 (cyan) and surrounding regions of uOmpA_171_ and the core and P1 domains of SurA. **(C)** Representative structural model of the third binding mode (o1s009), supported by fifteen crosslinks between segment 1 (pink) and segments 3, 4, and 5 (orange) primarily to the core domain of SurA. **(D)** A representative structure of two SurA protomers bound to uOmpA_171_ (o2s006), wherein one copy of SurA binds segment 1, and another copy binds segments 3, 4, and 5.

As described above, we constructed (SurA)_*n*_•uOmpA_171_ models with higher-order stoichiometries (where *n* = {2, 3, 4}). These models provide an additional explanation for the complex DSBU crosslinking pattern between uOmpA_171_ and SurA. Figure 4D shows how a single uOmpA_171_ can distribute these binding modes across more than one copy of SurA instead of threading itself through a single copy of SurA multiple times – accommodating the same clusters of crosslinks. Moreover, inclusion of models with higher order stoichiometries increases coverage to 89% of identified DSBU crosslinks (Supplementary Data 6).

Finally, as a critical control to confirm the SurA groove as the primary binding site for uOMP substrates, we performed DSBU crosslinking experiments between uOmpA_171_ and the “locked-closed” SurA variant (P61C/A218C) previously described by Silhavy and co-workers (38).

In this variant the open conformation of SurA is inaccessible, and cells show increased sensitivity to viability envelope stressors *in vivo*. We observed a drastic reduction in the number of “locked-closed” SurA–uOmpA_171_ interprotein crosslinks compared to WT SurA (7 vs 46; see SI Appendix, Figure S12, Supplementary Data 3 & 5). This finding implies that the dense crosslinking patterns revealed by XL-MS depend on the formation of the SurA groove, and not from contacts made outside of the groove.

### SANS validates and quantifies XL-MS based binding modes

To test which SurA•uOmpA_171_ models are representative of the conformational ensemble in solution, we compared them to an independent SANS profile of SurA_105,*p*AF_-uOmpA_171_ collected in 0% D_2_O (see SI Appendix, Figure S14). In this particular condition, protonated SurA and perdeuterated uOmpA_171_ contribute equally to scattering (see SI Appendix, Figure S6). We calculated the expected scattering profile that would arise under these experimental conditions for each of the 40 SurA•uOmpA_171_ models using the software package, SASSIE (70). In addition to the models of the SurA•uOmpA_171_ complex, we included multiple models of apo-SurA in varying conformations as we were unable to completely purify the crosslinked complex (see SI Appendix, Figure S15). In agreement with the DSBU crosslinking, we found that no single model, or pairs of models, recapitulated the experimental scattering profile (reduced chi-square < 1.05) (71).

Linear combinations of the scattering profiles from three structural models were able to describe the SANS dataset. In total, we sampled over 1 million combinations of triplets of models and found 35 combinations whose simulated SANS profiles produced reduced chi-square values less than 1.05 with respect to the experimentally observed 0% D_2_O SANS profile. Each accepted triplet contained at least one model of non-crosslinked SurA and one SurA•uOmpA_171_ complex (see SI Appendix, Table S7). The three models most likely to be included in an accepted combination arise from the three distinct binding modes defined by XL-MS, and are shown in Figure 4A-C with their associated crosslinks. In addition, the linear combinations identify several models with higher-order stoichiometries as depicted in Figure 4D.

The sparse ensemble illustrating the main structural features identified by our experiments is depicted in Figure 5 by an overlay of SurA•uOmpA_171_ models (population of each model in the ensemble are listed in Supplementary Appendix, Table S8). We note that this collection of conformations captures all known experimental data on SurA•uOMP complexes: including uOMP binding to the SurA groove (Figure 1), the Hill coefficient that is slightly greater than 1, the expansion of uOMPs by SurA (Figure 2), pXL-MS identification of specific binding segments (Figure 3), and the independent validation of binding modes by SANS and XL-MS (Figure 4 and 5) (22, 37, 38, 49). This sparse ensemble of models of the SurA•uOmpA_171_ complex provides a chemically reasonable and minimalist set of structures that could exist in the full conformational ensemble of the complex in solution. Due to limitations in the resolution of the data, additional conformations that fit the data likely exist.

**Figure 5.**
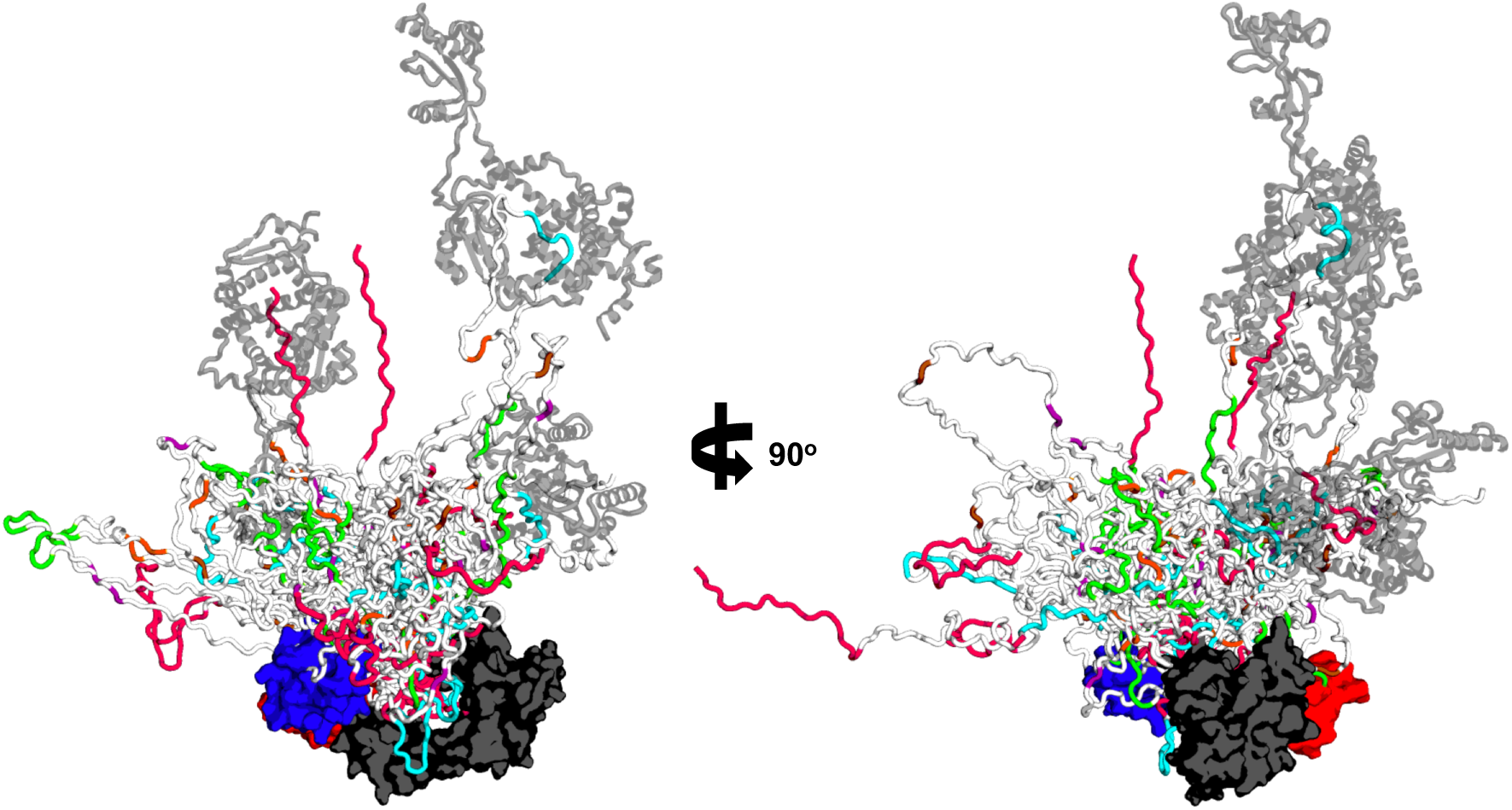
SurA•uOmpA_171_ ensemble as defined by experimental restraints reveals uOmpA_171_ conformational landscape. The 21 SurA•uOmpA_171_ structural models that were part of triplets of linearly weighted models that fit the 0% D_2_O SANS data are overlaid aligned to SurA. SurA is in the open conformation and shown with a surface representation with domains colored as in Figure 1A. uOmpA_171_ models are shown with a cartoon representation and have SurA-binding segments, as defined by XL-MS, colored as in Figure 3. Higher order stoichiometries are found in the ensemble, with additional copies of SurA shown as transparent, gray cartoons. The diversity of uOmpA_171_ conformations that are shown in this ensemble highlight the conformational dynamics accessible to client uOMPs when bound to SurA.

## Discussion

The periplasmic chaperone network is integral for *E. coli* outer membrane protein biogenesis. SurA plays several important roles in the uOMP biogenesis pathway: it must (i) recognize uOMP clients before they aggregate; (ii) maintain them in a soluble form in the periplasm; and (iii) mediate a hand-off to the BAM complex. In this study we have utilized an integrative/hybrid structural biology approach that combines multiple crosslinking and scattering techniques to generate restraints used to build a representative ensemble. This ensemble captures key structural features of SurA in complex with its client uOmpA_171_.

The model-independent Guinier analysis of the contrast-matched SANS experiments reveal that SurA performs its functions by dramatically expanding the SurA-uOMP complex in a mechanism reminiscent of trigger factor, a structural homolog to SurA (72). We observe a primary uOMP interaction in the SurA groove, located between the core and P1 domains which recapitulates recently published findings (37). The remainder of the unfolded uOmpA_171_ chain that is not occupying the groove must assume an elongated conformation to be consistent with these SANS data. In contrast, a covalently “locked-closed” variant of SurA is unable to efficiently interact with uOmpA_171_, consistent with the reduced functionality of this variant *in vivo*. All together these results suggest uOMP binding to the SurA groove results in uOMP expansion relative to its compact native state, as exemplified in Figures 3, 4, and 5.

This expansion of uOMPs is distinctive from a recently published model of the SurA-uOmpX complex built from crosslinking data alone (37). In that model, a single SurA completely encapsulates a globular uOMP, and this structural interpretation is incongruous with our experimental findings as the present SANS data demonstrate that the uOmpA_171_ must instead be expanded. Although the reported crosslinking is consistent with our data, our approach also capitalized upon the usefulness of a hydrodynamic view of highly dynamic structures. Instead, our data for the SurA-uOmpA_171_ complex are more consistent with a dynamic conformational ensemble proposed for the SurA-FhuA complex based on NMR (42). The extent of expansion of unfolded FhuA is unresolved, however, because the hydrodynamic properties of this system have not yet been established.

Unfolded OMP expansion mediated by a single SurA binding event is aided by the ability of additional copies of SurA to interact with distal binding segments on the uOMP (as shown in Figures 3B and 4D). Expansion of uOMPs would help avoid steric clashes between different copies of SurA simultaneously interacting with an uOMP client. For the small, eight-stranded uOMPs investigated here, higher-order stoichiometries represent minor populations in the ensemble. It is reasonable to speculate that the length of a client uOMP could dictate the binding stoichiometry of the SurA•uOMP complex as larger clients would ostensibly contain a greater number of SurA-binding segments. In accordance with this idea, gel filtration data suggest that the stoichiometry for the SurA-FhuA complex may be closer to 2:1 because FhuA is a much larger outer membrane protein (22 β-strands as compared to 8 for OmpA) (31, 42). Higher-order stoichiometries could also be enhanced in the crowded periplasm, where the excluded volume effect increases protein-protein interactions (73).

The initial finding of uOmpA_171_ expansion was surprising, especially because the other major periplasmic chaperone Skp encapsulates uOMPs in a collapsed state reminiscent of their intrinsic, unfolded conformation (42, 60). Given SurA interacts with uOMPs primarily through its groove, the persistent global expansion of the remainder of the client uOMP at first glance appears puzzling. We propose a kinetic trapping mechanism wherein binding and release of uOMP segments are fast relative to the collapse of the uOMP to its intrinsic molten globule state. Indeed, kinetic partitioning is a dominant organizing feature of the uOMP biogenesis chaperone network, and the interaction between SurA and uOMP happens on a very fast time scale (18, 43, 74). Rates of unfolded uOMP collapse are not known but may be relatively slow given the low overall hydrophobicity of transmembrane β-barrel primary sequences. Such a difference in the rates of uOMP intrinsic collapse versus expansion by SurA binding would provide a way to retain uOMPs in an expanded state through transient repeated associations with SurA protomers. This mechanism has the advantage of limiting the amount of SurA required to solubilize a client thereby maximizing the reservoir of free SurA in the periplasm.

Figure 6 highlights several biological implications of the features of our structural ensemble. We hypothesize that these SurA properties could explain its multifaceted roles in OMP biogenesis. Firstly, we expect that expansion of uOMPs would decrease unproductive intraprotein interactions and maintain the chain in a folding-competent, unfolded conformation (Figure 6B). In this respect, SurA performs an orthogonal role to the other chaperones in the uOMP biogenesis network, which form cages around uOMPs to decrease interprotein interactions. Expansion may also mediate the formation of transient hetero-chaperone complexes in the periplasm, where more than one chaperones are simultaneously bound to an uOMP (Figure 6C and D). As uOMPs are expected to undergo hundreds individual binding and dissociation events while in the periplasm, a SurA-mediated hand-off of expanded uOMPs between chaperones would allow for these events to occur while keeping the population of aggregation-prone, unbound uOMP low (18).

**Figure 6.**
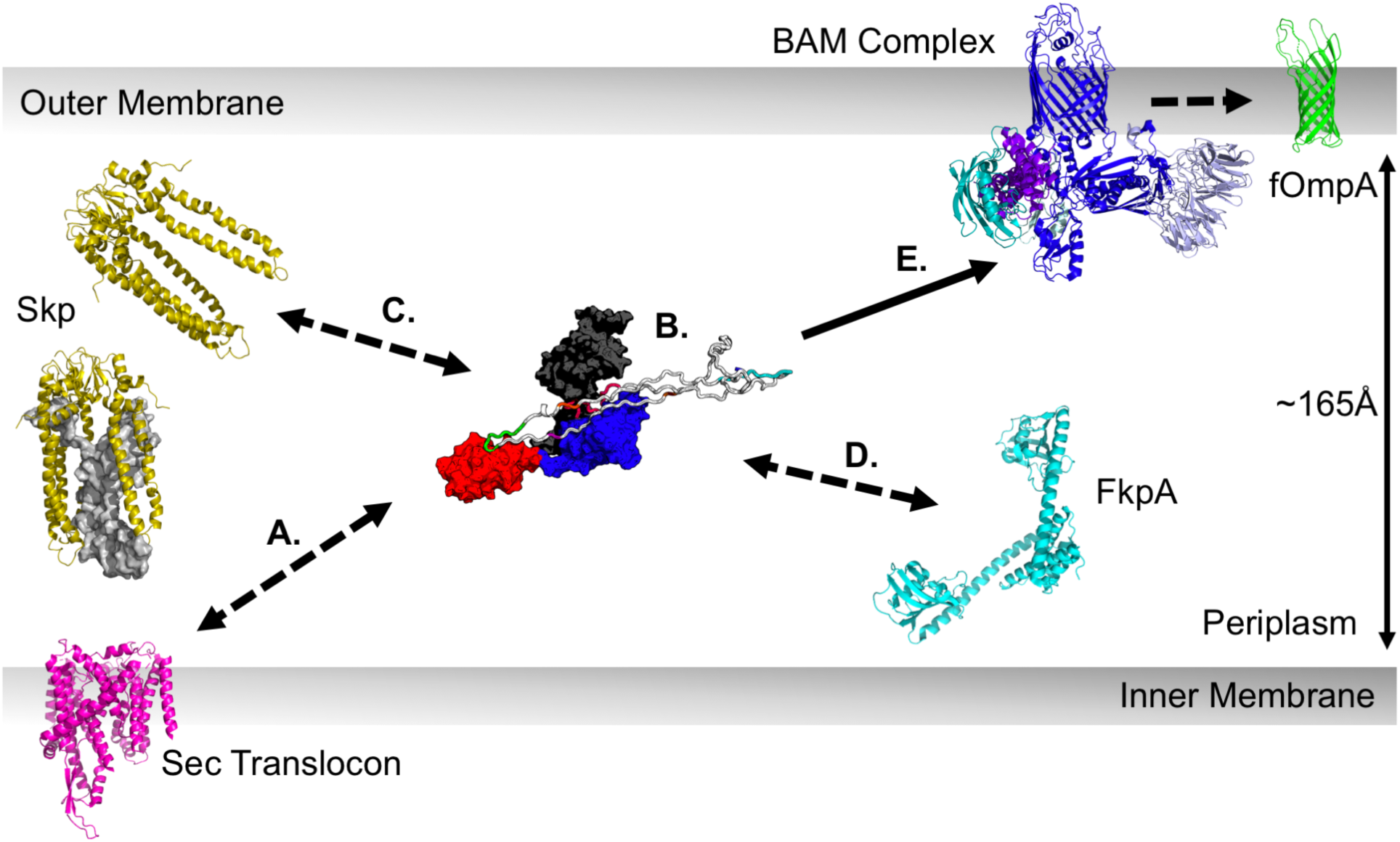
Implications of uOMP expansion in the periplasm. uOMPs are post-translationally secreted through the Sec translocon (magenta), N-to-C terminally. **(A)** The emerging uOMP N-terminus in the periplasm may be recognized by SurA. **(B)** After complete translocation into the periplasm, one or more SurA protomers bind specific segments on uOMP clients, solubilizing the uOMP in expanded conformations nearly the width of the periplasm. The expanded uOMP may also be able to form heterocomplexes with other chaperones in the OMP biogenesis pathway: Skp **(C)** and FkpA **(D)**. The size of uOMPs bound to SurA is approximately double the size of uOMPs bound to Skp. Unfolded OmpA_171_ bound to Skp is shown as a gray surface representation in the Skp trimer located proximal to the translocon. **(E)** The extended, unbound C-terminal region of the SurA-bound uOMP is positioned to encounter the BAM complex, which recognizes the OMP β-signal and catalyzes uOMP folding into the outer membrane.

An intriguing outcome of our structural ensemble is the finding that SurA-binding segments of client uOMPs are located toward the N-terminus (Figure 3). Indeed, the three binding modes found for the SurA•uOmpA_171_ interaction are mediated by the two most N-terminal binding segments on uOmpA_171_. This suggests the possibility of an early encounter between SurA and client uOMPs in the periplasm (Figure 6A). Accordingly, a recent, low resolution cryo-EM model places SurA near the translocon where it is well positioned to bind emerging uOMP segments (75).

Conversely, there was a conspicuous absence of robust crosslinking for both client OMPs near their C-termini. The apparent lack of SurA interaction sites in this uOMP region leaves the β-signal (Aro-X-Aro) free to interact with other members of the uOMP biogenesis pathway. The β-signal has been shown to play an important role in mediating efficient catalysis of uOMP folding by BAM both *in vivo* and *in vitro* (76–78). As SurA is the only periplasmic chaperone that promotes the interaction between uOMPs and BAM, our data support a mechanistic hypothesis in which this region of uOMPs is free and flexible and effectively ‘cast’ outward in a mechanism reminiscent of fly fishing to catch the BAM complex (Figure 6E) (13, 79).

Even if it is transient, the formation of a SurA•uOMP•BAM ternary complex is enticing because it brings the uOMP close to both the BAM complex and the disrupted adjacent membrane, both of which accelerate uOMP folding (9, 80). The BAM complex has been proposed to template and insert uOMP β-hairpins as individual foldamers of OMPs. Notably, the SurA groove is large enough to accommodate a β-hairpin, which could potentially favor the pre-formation of this key structural elements in a nascent uOMP. Moreover, this SurA-mediated β-hairpin formation mechanism could be easily adapted to larger clients with more transmembrane strands (and probably more SurA-binding segments) given the modular nature of the β-hairpin unit.

In this work we highlight the utility and complementarity of photo- and chemical crosslinking, neutron scattering, and mass spectrometry applied together. This combined approach was crucial because unfolded proteins present many challenges to conventional structural techniques due of their absence of regular secondary structure, their high conformational flexibility, and their propensity to aggregate. Our results illuminate a sparse ensemble of models that captures the key structural features defining how SurA promotes outer membrane protein biogenesis.

## Materials and Methods

### SurA Expression and Purification

The **SI Appendix** describes creation, expression, and purification of all SurA constructs used in this study.

### *p*AF Crosslinking

25 µmols L^-1^ (µM) of each SurA_*p*AF_ variant was mixed with 5 µM uOMP (20 mmoles L^-1^ (mM) Tris, 1 mol L^-1^ (M) urea, pH = 8.0. uOMPs expressed and purified as described previously (81). We chose these conditions because both SurA and uOmpA_171_ are monomeric and soluble at the listed protein and urea concentrations (39, 51). Mixtures were then irradiated with UV light (wavelength, λ= 254 nm) for 5 minutes using a Spectroline MiniMax UV Lamp (Fisher #11-992-662). Aliquots were taken for SDS-PAGE analysis both pre- and post-exposure to UV light. These samples were subjected to electrophoresis using a 4-20% gradient precast gel (Mini-PROTEAN TGX, Bio-Rad) at a constant voltage of 200 V for 35 minutes at room-temperature.

Using ImageJ, densitometry analysis on the loss of density of the uOmpA_171_ band was utilized to quantitate crosslinking efficiency. Crosslinking efficiency values were corrected for the amount of uOmpA_171_ band lost (∼20%) when mixed with WT SurA (not containing *p*AF). A representative SDS-PAGE gel for each SurA_*p*AF_ variant and uOmpA_171_ is shown in the SI Appendix, Figure S2. This same protocol was utilized to assess the crosslinking of SurA_*p*AF_ variants to uOmpX and uOmpLA.

### SANS on Protonated-SurA/Perdeuterated-uOmpA_171_ Complex

SurA_105,pAF_ was crosslinked to deuterated-uOmpA_171_ as described above. Perdeuterated OmpA_171_ growth, expression, purification, and characterization are detailed in the SI Appendix. This complex was further purified via size exclusion chromatography (GE Superdex-200 10/300 GL; flowrate = 0.6 mL/min) in 20 mM Tris, 200 mM NaCl, pH 8.0 (GF buffer), and buffer exchanged into either 0 % or 30 % D_2_O for SANS experiments (same buffer components as SEC) (see SI Appendix, Figure S15). We made three attempts to also collect scattering profiles in 80% and 98% D_2_O of this complex but the I(0) values from Guinier fitting indicated that these samples contained aggregates. It is known that increased buffer concentrations of D_2_O may promote self-association and aggregation of particularly hydrophobic proteins (82).

All scattering experiments were collected at the National Institute of Standards and Technology Center for Neutron Research (Gaithersburg, MD) as previously described(60). More information on SANS data collection and analysis are found in the **SI Appendix**.

### XL-MS of SurA-uOMP complexes

The **SI Appendix** describes the methods used to perform and analyze all XL-MS experiments, including: SurA_*p*AF_-uOMP photo-crosslinked complexes, SurA-uOMP DSBU crosslinked complexes, and “locked closed” SurA-uOmpA_171_ DSBU crosslinking complexes. This includes *p*AF and DSBU methods and data analysis protocols.

### Structural models of SurA

Models of SurA were constructed based on crystal structures 1M5Y and 2PV3 (Table S2). In the 1M5Y structure, the core and P1 domains are close together but the P2 domain is extended. In the 2PV3 structure, the P1 domain is moved away from the core domain and rotated relative to 1M5Y but the P2 domain is missing. Residue segments 20-34, 165-171, 387-394, and 428-430 were built into 1M5Y and six histidine residues were added to the C-terminus using Modeler to create the “P1 closed” form of SurA (83). For the “P2 closed” form of SurA, PyMOL was used to build the P2 domain from the P1 closed form into 2PV3 (84). The P2 domain was then moved into position against the groove formed by the core and P1 domains using NAMD as described below. The “open” SurA model has the core-P1 relative orientation from 2PV3 and the core-P2 relative orientation from 1M5Y. Domains were oriented in PyMOL and linkage conformations were normalized using NAMD as described below. The “Collapsed” SurA model has the core-P1 relative orientation from 1M5Y and the P2 domain was moved into position against the core using NAMD as described below.

The P2 closed, open, and collapsed SurA models, initially constructed using PyMOL, were further manipulated to position the domains, remove Van der Waals clash and relax unstructured linkage segments using NAMD with generalized Born implicit solvent electrostatics in the CHARMM22 force field (85). Domains were positioned with targeted distance restraints as implemented in the collective variable module in NAMD (86, 87). Typically, a harmonic potential was placed on the distance between the centers of mass of two groups of CA atoms with a force constant of 1.0 kcal/mol and the force was applied for 50,000 to 150,000 steps. This *in vacuo* molecular relaxation and manipulation was carried out after 200 steps of energy minimization, with implicit solvent alpha cutoff = 12.0 Å, [ion] = 0.3 M, non-bonded cutoff = 14.0, switching starting at 13.0, and 2 fs time step. Langevin dynamics was used with a damping coefficient of 1 for temperature control (NVT). The domain-domain distances of SurA were monitored during simulation and a structure was saved when target distances were obtained.

### Structural models of the SurA•uOmpA_171_ complex

Four extended uOmpA_171_ segments (residues 2-21, 54-73, 84-104, 112-132) that contain the six SurA-binding segments were independently submitted, along with the open SurA structural model to the protein-protein docking web server HADDOCK (66). These sequence segments were chosen to include those residues that were found to repeatedly crosslink to the high efficiency SurA_pAF_ variants. Active and passive residues for HADDOCK were chosen from SurA groove (see SI Appendix, Figure S9).

These docked oligopeptides were inspected using molecular graphics to obtain target distances for docking the full length, unexpanded uOmpA_171_ models (s = 1.65) to the open form of SurA. Docking was accomplished in NAMD using the target distances from HADDOCK peptide docking as distance restraints in the collective variables module of NAMD as described above.

The uOmpA-open SurA models were further manipulated to increase the uOmpA D_Max_ to the target of 150 Å that was determined by P(r) analysis of the SANS data. These expansions were accomplished using distance restraints and the collective variable module in NAMD. Short segments of each bound uOmpA that were furthest apart were identified and the two groups of respective CA atoms were forced to a distance of ∼150Å with a harmonic potential as described above. A second open SurA model was then docked to exposed, known binding segments (see SI Appendix, Figure S11) of the extended uOmpA. In three cases, a third open SurA was docked to remaining exposed known binding segments. One extended polypeptide of uOmpA was generated with a D_Max_ ∼250 Å and four open SurA models were docked to the four main segments on OmpA that displayed high efficiency cross-linking.

In all, twenty-three models containing one docked SurA, thirteen models containing two docked SurA, three models containing three SurA, and one model containing four SurA were built. Physical dimensions of these models are listed in Table S6. Values for *R*_G_ and *D*_Max_ were calculated using HullRad(63). All models contained CHARMM hydrogens and were used to calculate predicted SANS profiles using the SasCalc server.(70)

Methods and information regarding the comparison of structural models to the 0% D_2_O SANS profiles performed to generate the ensemble of structures shown in Figure 5 are found in detail in the **SI Appendix**. Models included in the sparse conformational ensemble of SurA-uOmpA171 can be found at: https://github.com/KarenGFleming/SurAuOmpA.

## Data access statement

All raw data and detailed protocols including gel images, SANS profiles, and model creation protocols, are available upon request. The SI Appendix includes detailed protocols and raw data associated with XL-MS experiments.

## Abbreviations

SurA•uOMP: non-crosslinked protein-protein complex
SurA-uOMP: crosslinked protein-protein complex
uOMP: unfolded outer membrane protein
BAM: beta barrel assembly machinery
*p*AF: para-azido phenylalanine
pXL-MS: photo-crosslinking mass spectrometry
SASD: solvent accessible surface distance

## Acknowledgements

We thank the Johns Hopkins University Biomolecular NMR Center and the Center for Molecular Biophysics for providing facilities and resources. Access to NGB30 SANS was provided by the Center for High Resolution Neutron Scattering, a partnership between the National Institute of Standards and Technology and the National Science Foundation under Agreement No. DMR-1508249. We acknowledge the support of the National Institute of Standards and Technology, U.S. Department of Commerce, in providing the neutron research facilities used in this work. Certain commercial equipment, instruments, or materials (or suppliers, or software, …) are identified in this paper to foster understanding. Such identification does not imply recommendation or endorsement by the National Institute of Standards and Technology, nor does it imply that the materials or equipment identified are necessarily the best available for the purpose. This work benefitted from CCP-SAS software developed through a joint EPSRC (EP/K039121/1) and NSF (CHE-1265821) grant. This work was supported by National Science Foundation (NSF) MCB1412108 and NIH R01 GM079440 grants (to K.G.F.). D.C.M, A.M.P., T.D. and H.J.L. were supported by NIH training grant T32 GM008403. A.M.P. was supported by the NSF grant DGE 1232825. The authors thank lab members for helpful discussions.

