## Supplemental Information for "SurA is a “Groove-y” Chaperone That Expands Unfolded Outer Membrane Proteins"

**This PDF file includes:**

Supplementary text
Figures S1 to S16
Tables S1 to S9
Legends for Datasets S1 to S6
SI References

**Other supplementary materials for this manuscript include the following:**

Datasets S1 to S6

### Supplemental Methods

***SurA Expression and Purification*** **-** We introduced the gene of E. *coli* SurA lacking the signal sequence into the pET28b vector between the NdeI and BamHI restriction sites with a C-terminal 6-Histidine tag. The library of SurA$_{pAF}$ variants were created by incorporating an Amber stop codon (TAG) into 32 individual positions in the gene (QB3 Berkeley Microlab) (Figure S1) (Certain commercial equipment, instruments, material, suppliers, or software are identified in this paper to foster understanding. Such identification does not imply recommendation or endorsement by the National Institute of Standards and Technology, nor does it imply that the materials or equipment identified are necessarily the best available for the purpose.). Surface exposed sites were chosen using the 1M5Y crystal structure of apo-SurA. The "locked-closed" SurA variant was created by cloning in two point-mutations (P61C/A218C) using InFusion Cloning (Takara). Plasmids were transformed into HMS E. *coli* cells. For *p*AF incorporation, HMS cells also harbored the pDule2 plasmid which encodes the amber-suppressor tRNA$_{Tyr(CUA)}$ from *Methanocaldococcus jannaschii*, and its cognate aminoacyl-tRNA synthetase (gracious gift from the Sondermann lab).

Following an overnight growth, 500 mL Terrific Broth cultures were supplemented with 50 μg mL$^{-1}$ kanamycin (10 μg mL$^{-1}$ streptomycin was also added to pDule2 containing cultures) and induced at O.D.$_{600}$ = 0.6-0.8 with IPTG (final concentration 0.1 mmol L$^{-1}$ (mM)). *p*AF was added to a final concentration of 1 mM at the time of induction for SurA$_{pAF}$ variant growths. Cells were harvested after growth overnight (5000 rpm for 30 min, 4°C) in a Beckman J2-MI centrifuge (JA-10 rotor) and pellets were frozen and stored at -20°C.

For purification, cell pellets were solubilized in Buffer A (20 mM sodium phosphate, 500 mM NaCl, 20 mM imidazole, pH 8.0; one Pierce EDTA-free protease inhibitor tablet added, (Thermo Prod # 88266)) and subsequently lysed using an Avestin Emulsiflex homogenizer. Lysate (supernatant) was harvested via ultracentrifugation (5000 rpm for 30 min, 4°C), then filtered (0.22 μm pore size) and purified using a Ni-NTA Sepharose High Performance bench-top column. After loading onto the column, the sample was washed with 20mL of Buffer A containing 6M urea. Samples were refolded on the column and eluted in Buffer A containing 300 mM imidazole. Purified SurA was then dialyzed overnight (10 kDa MWCO Snakeskin dialysis tubing, Thermo Prod #68100) into 20 mM Tris, pH 8.0, concentrated using a 30 kDa MWCO Amicon spin concentrator (Millipore) and stored at -

20°C. Stock concentrations were determined with the theoretical extinction coefficient of 29450 $M^{-1}$ $cm^{-1}$ (1).

***Perdeuterated OMP Expression, Purification, and Characterization.*** Deuterium was incorporated into the $uOmpA_{171}$ protein as previously described(2). Briefly, we expressed $uOmpA_{171}$ to inclusion bodies in minimal M9 growth media containing $D_2O$ and deuterated-glucose. Inclusion bodies were isolated and stored in -20 °C. Prior to use, inclusion bodies were thawed and solubilized in 8 M Urea, 20 mM Tris.

To determine the extent of OMP perdeuteration, which is a required parameter for SANS contrast calculations, we utilized one-dimensional proton NMR. We collected 1D $^1H$ spectra on both protonated and deuterated $uOmpA_{171}$ (50 μM) in 8 M Urea, 20 mM Tris, 10 % $D_2O$ at 35 °C, on a 600 MHz Bruker Avance II spectrometer (Figure S16). Water suppression was achieved using a flipback-watergate sequence and a buffer purging pulse was included to minimize the large urea peak. Each spectrum was collected using 128 scans, a recycle delay of 1.5 s, and acquisition time of 150 milliseconds per Free Induction Decay (FID) scan. Data was processed and analyzed using TopSpin 2.1. The spectra were aligned using the amide resonance peaks, and the baseline of the methyl peaks was corrected using a $5^{th}$ order polynomial. After baseline correction, the methyl peak volumes were integrated using TopSpin, and the integrated intensities of the methyl peaks in both the protonated and deuterated samples were compared. The loss of intensity in the methyl peaks between the two samples was used to estimate the deuteration level of deut-$uOmpA_{171}$. OMP deuteration was estimated to be 80 %.

***SANS Data Analysis.*** For analysis of SANS datasets, we utilized the Guinier approximation to obtain two fit parameters: the macromolecule $R_G$ (Å) and the forward scattering intensity at q = 0 (i.e., I(0) in $cm^{-1}$). This approximation estimates the intensity in low q regions as follows:

$$I(q) \approx I(0)\exp\left[-\left(\frac{1}{3}\right) R_G^2 q^2\right] \quad \text{(Equation S1.)}$$

$$\ln[I(q)] \approx \ln[I(0)] - \frac{1}{3} R_G^2 q^2 \quad \text{(Equation S2.)}$$

Therefore, linear regression (i.e., Figures 2 and S14) of ln[(I(q)] vs. $q^2$ yields information in the slope (i.e. $R_G^2$) and the intercept (i.e., I(0)). I(0) was also calculated using the Contrast Calculator module (3) in the web version of the SASSIE software developed at

NIST (4). Our complexes contain components with different deuteration levels and contrast, which is take into account where:

$$I(0) = \frac{CM}{N_A}(\sum_i f_i \Delta\rho_i \, \bar{v}_i)^2 \qquad \text{(Equation S3.)}$$

where C indicates the protein concentration in g $mL^{-1}$, $N_A$ is Avogadro's number, $\Delta\rho$ is the component contrast, and $\bar{v}$ is the component partial specific volume mL $g^{-1}$, $f_i$ is $M_i/M$, M is molecular weight in Da, and the summation is over the two components of the complex. This sum includes three terms: one for protonated SurA, one for perdeuterated $uOmpA_{171}$, and a cross-term that originates from inter-protein interactions. At 30 % $D_2O$ using this rigorous calculation method, SurA, $uOmpA_{171}$, and the cross-term contribute to the total I(0): 9 %, 48 %, and 43 % respectively, the main contribution from the cross-term coming from the $\Delta\rho$ value of $uOmpA_{171}$.

To better understand the contribution to the total scattering of SurA and $uOmpA_{171}$ in the absence of the cross-term, we approximated the contribution of these components to the total scattering using the following simplified formula:

$$I(0) = \frac{CM}{N_A}\sum_i (f_i \Delta\rho_i \bar{v}_i)^2 \qquad \text{(Equation S4.)}$$

At 30 % $D_2O$ using this simplified calculation method, SurA and $uOmpA_{171}$ contribute 16 % and 84 % to the total I(0), respectively.

For each Guinier fit, we compared the fit value of I(0) to the calculated value for each experiment (Table S3).

In addition to Guinier fitting, we obtained $R_G$ and I(0) values from distance distribution functions, P(r) vs. r (Table S4). These fits were completed using autoGNOM(5) and a range of 0.011 $A^{-1}$ to approximately 0.2 $A^{-1}$. Fit values reported in Table S4 were obtained using the specified $D_{max}$ values from autoGNOM. The P(r) vs. r curve is shown in Figure 2. Exploration of a range of $D_{max}$ values near the specified values did not result in a change in the shape of P(r) versus r, except for small changes in the region near $D_{max}$, as shown in Supplemental Table S4.

***Evaluation for Agreement between SANS Data and Form Factors***

We evaluated agreement between the SANS scattering profile collected in the 0% $D_2O$ condition and the scattering form factors from models using the reduced $\chi^2$ according to Equation S4 (6):

$$\chi^2 = \frac{1}{N-M}\sum_1^N \left[\frac{I(q)_{obs} - I(q)_{calc}}{\sigma_{obs}}\right]^2 \qquad \text{(Equation S5.)}$$

where $N$ equals the number of data points, $I(q)_{obs}$ and $I(q)_{calc}$ are the experimental and calculated intensity values, respectively, at each point $q$ and $\sigma_{obs}$ is the error on the experimental measurement at each point. A good fit is defined as $\chi^2 < 1.05$.

***Comparison of Structural Models Experimental 0% $D_2O$ SANS Profiles***

The SasCalc module in SASSIE was used to calculate SANS form factor profiles $P(q)_{calc}$ for all models (4, 7). Normalized form factor curves were obtained by dividing $P(q)_{calc}$ by the I(0) for a given data set. SasCalc form factors were evaluated for their ability to describe the corresponding, experimental SANS curves using the equation for the reduced $\chi^2$ (equation S4) as recommended by Trewhella and colleagues (6). As expected for an ensemble, a good fit was not obtained by a single structural model. Therefore, fractionally weighted, linear sums of SasCalc form factor curves were calculated. These included the appropriate weighting for different scattering contrast values between SurA and $uOmpA_{171}$. The equations for basis set addition for cases where the components in solution do not possess uniform scattering contrast are described next. For a linear addition of two basis form factors, the composite scattering curve predicted for a structural model, $I(q)_{calc}$, is a function of the contrast and weight contributions of each component as follows:

$$I(q)_{calc} = \left[W_i P(q)_{i,calc} + W_j P(q)_{j,calc}\right] \qquad \text{(Equation S6.)}$$

where the $W_i$ terms correspond to the fractional contrast weighting terms simulated from a combination of weight fractions $(F_i)$. The weighting terms were obtained from contrast factors calculated for each component as follows:

$$W_i = \frac{C_i}{C_i + C_j} \qquad \text{(Equation S7.)}$$

where $W_i + W_j = 1$, and the $C$ variable in these equations for each component equals:

$$C_i = \frac{c_{Tot} F_i m_i \left(\sum_a^N f_a \Delta\rho_a \bar{v}_a\right)^2}{N_A} \qquad \text{(Equation S8.)}$$

where $c_{Tot}$, N, $m$, $\Delta\rho$, and $\bar{v}$ are the weight concentration, molar mass (g mol$^{-1}$), contrast values (Table S9), and partial specific volume (ml g$^{-1}$) for each species, respectively $f_a$ in a complex is the mole fraction of the $a^{th}$ component within that complex; and $\sum_1^A f_a = 1$ where $A$ is the number of components with any given complex. For normalized data the $c_{Tot}$ term can be dropped because its value has no effect on the $W$ values. For example, the scattering equation for a pair of component curves containing a one-to-one complex and free SurA in cases of nonuniform scattering equals:

$$C_{SurA} = \frac{F_i m_{SurA}(\Delta\rho_{SurA}\bar{v}_{SurA})^2}{N_A} \quad \text{(Equation S9.)}$$

and

$$C_{Complex} = \frac{F_j m_{Complex}\left(f_{SurA_{105}}\Delta\rho_{SurA_{105}}\bar{v}_{SurA_{105}} + f_{OmpA}\Delta\rho_{OmpA}\bar{v}_{OmpA}\right)^2}{N_A}$$

(Equation S10.)

Although the $SurA_{pAF105}$-$uOmpA_{171}$ crosslinked sample formed a distinct 1:1 band as assessed by SDS-PAGE, the excess free SurA was not completely separated from this sample by subsequent gel filtration. Repeat experiments of mock sample preparation indicate that this non-covalently bound SurA is present at mole ratios of 0.3 to 1 and must therefore be taken into account in the SANS profile analysis. This technical issue turned out to be fortuitous because it allowed the population of low levels SurA-uOmpA171 to bind additional SurA protomers, thus populating complexes with multiple SurA molecules bound.

To simulate a wide range of linear combinations, we iterated through weight fractions at 1% increments and compared these to the 0% SANS curves using the reduced $\chi^2$ (equation S5 above) with M increased to 2 to account for the additional loss of degrees of freedom. No pairwise combination of form factors resulted in a good agreement with the data as evidenced by acceptable $\chi^2$ and appropriate mole ratios of non-crosslinked to crosslinked SurA.

We therefore extended these equations to three terms for triplets by the addition of a third term in equations, e.g.

$$I(q)_{calc} = \left[W_i P(q)_{i,calc} + W_j P(q)_{j,calc} + W_k P(q)_{k,calc}\right] \quad \text{(Equation S11.)}$$

and

$$W_i = \frac{C_i}{C_i + C_j + C_k} \quad \text{(Equation S12.)}$$

and

$$W_i + W_j + W_k = 1 \quad \text{(Equation S13.)}$$

and the terms are as described above and evaluating using the reduced $\chi^2$ equation above with M increased to 3 to account for the additional loss of degrees of freedom as compared to the paired case. Each weight fraction within a simulated triplet was incremented by 1% to achieve an exhaustive search of model combinations.

***XL-MS of Photo-crosslinked SurApAF-uOMP.*** Acetonitrile (ACN), Optima formic acid (FA), trifluoroacetic acid (TFA), Tris, and urea were obtained from Fisher Scientific (Hanover Park, IL, USA). LiChrosolv LC-MS grade water was obtained from EMD Millipore Corporation (Darmstadt, Germany). Pierce Trypsin protease (catalog number 90305) and Glu-C (catalog number 90054) protease were also obtained from Thermo Fisher.

Crosslinked samples comprising of 25 μmol $L^{-1}$ (μM) $SurA_{pAF}$ (with *p*AF at sites: 59, 94, 105, 120, 233, 245, 260, or 424), and 5 μM $OmpA_{171}$ or 5 μM OmpX were reconstituted in 20 mM Tris pH 8.0, 1 M urea, and crosslinked as described in the previous section (typically on a 50 μL scale, ca. 50 μg total scale of protein). Following crosslinking, solid urea was added to a final concentration of 2 M. Trypsin (1 μg/μL stock concentration) was then added (typically 1 μL) to the samples at a 1:50 enzyme/substrate ratio. The samples were digested overnight at 25° C, 700 rpm on a thermomixer.

For each protein complex, we analyzed peptides from a standard single-trypsin digest as well as from a double digest with trypsin then Glu-C in serial, and this was conducted in technical duplicate to generate four separate injections for each sample analyzed. For the latter sample, we added 20 mM Tris pH 8.0 to dilute the urea concentration to 0.8 M, whereupon Glu-C (1 μg/μL stock concentration) was added (typically 1 μL) to a 1:50 enzyme/substrate ratio. These samples were then digested overnight at 30°C, 700 rpm on a thermomixer.

Both singly-digested and double-digested peptides samples were acidified by addition of small volumes of TFA (~1 μL) to a final concentration of 1% (vol/vol). Samples were then diluted with 0.5% TFA to a final volume of 1 mL to facilitate loading into the cartridges.

Solid phase extraction was carried out using Sep-Pak C18 vacuum cartridges (Waters, Milford, MA, USA) according to the following protocol: Cartridges were first conditioned (1 mL 80% ACN, 0.5% TFA) and equilibrated (4x 1 mL 0.5% TFA), before loading the sample slowly under a diminished vacuum (ca. 1 mL/min). The columns were then washed (4x 1 mL 0.5% TFA), and peptides eluted by addition of 1 mL elution buffer (80% ACN, 0.5% TFA).

During elution, vacuum cartridges were suspended above 15 mL conical tubes, placed in a swing-bucket rotor (Eppendorf 5910R), and spun for 2 min at 350 *g*. Eluted peptides were transferred from Falcon tubes back into microfuge tubes and dried using a vacuum centrifuge (Eppendorf Vacufuge). Dried peptides were stored at -80˚C until analysis. For analysis, samples were vigorously resuspended in 0.1% FA in Optimal water to a final concentration of 1 mg/mL.

Chromatographic separation of digests was carried out on a Thermo UltiMate3000 UHPLC system with an Acclaim Pepmap RSLC, C18, 75 μm x 25 cm, 2μm, 100 Å column. Approximately 2 μg of protein was injected onto the column. The column temperature was maintained at 40ºC, and the flow rate was set to 0.300 μL/min for the duration of the run. Solvent A consisted of 0.1% FA in 2% ACN, 98% water, and solvent B consisted of 0.1% FA in ACN. After accumulation of peptides onto the trap column (Acclaim PepMap 100, C18, 75 μm x 2 cm, 3μm, 100 Å column) for 10 min (during which the column was held at 2% solvent B), peptides were resolved by switching the trap column to be in-line with the separating column, and applying a 100 min linear gradient from 2% B to 35% B. Subsequently, the gradient was increased from 35% B to 40% B over 25 minutes, and then increased again from 40% B to 90% B over 5 minutes. The column was then cleaned with a saw-tooth gradient to purge residual peptides between runs in a sequence.

A Thermo Q-Exactive HF-X Orbitrap mass spectrometer was used to analyze the eluting peptides. A full MS scan in positive ion mode was followed by ten data-dependent MS scans. The full MS scan was collected using a resolution of 120,000 (@ m/z 200), an AGC target of 3E6, a maximum injection time of 100 ms, and a scan range from 350 to 1500 m/z. The data-dependent scans were collected with a resolution of 15,000 (@ m/z 200), an AGC target of 2E5, a minimum AGC target of 8E3, a maximum injection time of 250 ms, and an isolation window of 2.0 m/z units. To dissociate precursors prior to their re-analysis by MS2, peptides were subjected to a stepped HCD with 22%, 25%, and 28% normalized collision energies. Fragments with charges of 1, 2, and >8 were excluded from analysis, and a dynamic exclusion window of 60.0 s was used for the data-dependent scans.

***XL-MS of Photo-crosslinked SurApAF Data Analysis.*** MS data were centroided and converted to the mzML file format using the msConvert application in the ProteoWizard Toolkit (8), and then analyzed for crosslinks using MeroX Version 2.0 (9). *p*AF was added to the amino acid list with a mass of 188.06981084 Da ($C_9H_8N_4O$). The photo-crosslink was added to the crosslink tab with composition of $-N_2$ (-28.006148 Da), a maximum Cα-Cα distance of 30 Å, specificity site 1 as *p*AF, and specificity site 2 as any amino acid. For tryptic digests, protease sites were allowed after arginine and lysine residues, with lysine blocked by proline as a cleavage site. For double digests, protease sites were allowed at arginine, aspartic acid, lysine, and glutamic acid, with lysine blocked by proline as a cleavage site. For both tryptic digests, a maximum of three missed cleavages was allowed (four was allowd for double digests). For modifications, a maximum of two oxidations of methionine was allowed. Searches were conducted using a FASTA file that consisted only of the uOMP in consideration and SurA. Otherwise, MeroX default parameters (for scoring and FDR calculation) were used.

Upon reviewing the output, a MeroX score of 50 was selected as the acceptance cutoff for crosslinked peptide-spectrum matches (PSMs). This corresponds to a FDR cutoff of <0.01; and in some cases to a far lower cut-off (Table S5). In numerous situations, a crosslink site to *p*AF could not be pinpointed down to a specific residue within a given peptide-spectrum match because of insufficient fragment ion data. In these situations, if several PSMs were available in which one provided more specific identification of a crosslink site and others included that site as part of larger nonresolvable region, we then merged the data to take advantage of the greater specificity when it was available. In Supplementary Data 1 and 2, the full list of crosslink sites associated with all PSMs are provided. Lines colored purple represent the most specific crosslink site assignable, and lines colored blue have lower resolution of the crosslink site but are consistent with a crosslink at a more specific site. The crosslink sites are compiled across all the $SurA_{pAF}$ variants for each uOMP in a tab labeled 'compiled_ Omp_SurApafs.'

***XL-MS of DSBU-crosslinked SurA-uOMP.*** A 50 μL solution comprised of 20 μM WT SurA combined with 20 μM of either $uOmpA_{171}$, uOmpX, or $uOmpA_{171}$ (P61C/A218C), was prepared in 20 mM $NaP_i$ pH 8.0, 1 M urea. Crosslinking was carried out by adding disuccinimidyl dibutyric urea (DSBU, ThermoFisher) from a 100 mM stock in DMSO to a final concentration of 1 mM. The sample was then mixed, incubated at room temperature with agitation for 30 min, and then quenched by addition of 1 M Tris pH 8.0 stock to a final Tris concentration of 100 mM. Following crosslinking, solid urea was added to a final concentration of 2 M. Trypsin (1 μg/μL stock, Pierce) was added to the sample (ca. 1–2 μL) to a 1:50 enzyme:substrate ratio. The samples were digested overnight at 25 °C, 700 rpm on a thermomixer. For samples prepared with a serial trypsin–GluC digest, the trypsinolysis reactions were diluted with 20 mL $NaP_i$ pH 8.0 to lower the final urea concentration to 0.8 M. Then 1–2 μL of GluC (1 μg/μL, Pierce) was added to a 1:50 enzyme:substrate ratio, and the samples were digested again overnight at 30˚C, 700 rpm on a thermomixer. Both single and double digests were then acidified with TFA to a final concentration of 1% (vol/vol), diluted with 0.5% TFA to a final volume of 1 mL, and then desalted by solid-phase C18 extraction columns, as described previously. Preparation of the sample and analysis by nanoLC-MS/MS was conducted identically to the samples generated by photo-crosslinking, as described previously.

**XL-MS of DSBU-crosslinked SurA Data Analysis.** MS data were centroided and converted to the mzML file format using the msConvert application in the ProteoWizard Toolkit,(8) and then analyzed for crosslinks using MeroX Version 2.0 using the software's standard settings for DSBU (9). Notably, MeroX uses a slightly expanded set of crosslink site specificities for DSBU: site 1 is restricted to be lysines and N-termini, whereas site 2 has an expanded specificity to also include serine, threonine, and tyrosine. Identified crosslinked peptides were filtered to an FDR of 1% and used if they had a MeroX score greater than 50 (which in most cases corresponded to an FDR well below 1%). Peptide spectrum matches (PSMs) were pooled between four separate injections (two replicates with trypsin only, two replicates with trypsin/Glu-C serial digest) to assemble a list of PSMs, given in Supplementary Data 3–5 under tabs labeled 'total.' We then merged together these datasets and removed redundancies to create condensed lists of crosslinks, also provided in Supplementary Data 3–5 under tabs labeled 'combined.' In several cases, the crosslink site could not be uniquely pinpointed, as occurs when numerous nucleophilic residues occur close to one another within a given peptide. These uncertainties in the position of the crosslink site are shown explicitly in the Supplementary Data tables.

To analyze these crosslinks further in light of the structural models of SurA•uOmp$A_{171}$, solvent-accessible Ca-Ca surface-distances (SASDs) between 85 pairs of DSBU crosslink sites between SurA and uOmp$A_{171}$ were calculated using using JWalk (upto a maximum SASD of 85 Å, grid size 1 Å). This expanded list of residue pairs was created by calculating all possible crosslinks that could be associated with a PSM with ambiguous linkage sites. For instance, if the PSM could determine the crosslink site on SurA to be position 50, but could not confidently determine if the crosslink site on uOmp$A_{171}$ was position 44 or 49 (corresponding to XL ID 1 in Supplementary Data 3), then we calculated the SASD both between SurA50–uOmpA44 and SurA50–uOmpA49. For assemblies with higher stoichiometry (i.e., $(SurA)_n$•$uOmpA_{171}$ with $n = \{2,3,4\}$), we calculated 170, 255, or 340 different SASDs because the identity of the SurA crosslink site could have come from any copy of SurA. To use the previous example of XL ID 1, the SASDs between $SurA_A$50–uOmpA44, $SurA_A$50–uOmpA49, $SurA_B$50–uOmpA44, and $SurA_B$50–uOmpA49 would all be calculated.

Next, for each structural model, a 'short-list' of the most likely crosslink sites was constructed by taking whichever of the possible linkages associated with a given XL ID admitted the smallest SASD, and discarding the others. To use the previous example, whichever of the four different SASDs was the lowest (and its associated crosslink sites) was used for XL ID 1. This procedure thereby provides for each structural model a set of 46 non-redundant SASDs. The SASDs were converted into harmonic scores using the following rule: all SASDs < 25 Å were awarded a perfect score of 100. Otherwise, SASDs were converted to scores using the function: score = $100 - 0.09\times(\text{SASD} - 25)^2$. Any score that was negative was converted to 0.0001. Any XL ID that did not have any SASD calculated by Jwalk was inferred to be greater than 85 Å, and was given a score of 0.0001. This scoring algorithm awards high scores to all crosslinks with SASDs ≤ 35 Å, moderate scores to crosslinks with $35 \leq \text{SASD} / \text{Å} \leq 45$ Å, and low scores (less than 50) to all crosslinks with SASDs greater than 48 Å. Crosslink distance cutoffs were chosen based off the well-established literature values of 25-35Å as acceptable values(10). The decay function to award moderate scores accounts for the flexibility of both SurA and uOmpA that cannot be captured in a single, static structure. All SASDs and scores can be found in Supplementary Data 6.

A matrix of scores was constructed for all 46 crosslinks in the context of 23 various SurA•$uOmpA_{171}$ structural models. Using the spectral biclustering algorithm with the 'log' method as implemented in scikit-learn with 20 biclusters (4 for structural models, 5 for crosslinks), the rows and columns of the score matrix were permuted to generate an organization that revealed distinct clusters, as shown in Figure S13 and discussed in the main text.

### Supplemental Figures

***Figure S1. SurA Sequence with pAF sites highlighted.***

```
 20 MAPQVVDKVAAVVNNGVVLESDVDGLMQSVKLNAAQARQQLPDDATLRHQIMERLIMDQI
 80 ILQMGQKMGVKISDEQLDQAIANIAKQNNMTLDQMRSRLAYDGLNYNTYRNQIRKEMIIS
140 EVRNNEVRRRITILPQEVESLAQQVGNQNDASTELNLSHILIPLPENPTSDQVNEAESQA
200 RAIVDQARNGADFGKLAIAHSADQQALNGGQMGWGRIQELPGIFAQALSTAKKGDIVGPI
260 RSGVGFHILKVNDLRGESKNISVTEVHARHILLKPSPIMTDEQARVKLEQIAADIKSGKT
320 TFAAAAKEFSQDPGSANQGGDLGWATPDIFDPAFRDALTRLNKGQMSAPVHSSFGWHLIE
380 LLDTRNVDKTDAAQKDRAYRMLMNRKFSEEAASWMQEQRASAYVKILSNLEHHHHHH
```

The sequence of SurA is shown, with the residues replaced by para-azido phenylalanine (*p*AF) highlighted in magenta. These sites were chosen on the basis that they are surface exposed in the monomeric crystal structure of SurA (PDB: 1M5Y) and are polar in nature to minimize the effects of pAF incorporation on protein structure and function.

***Figure S2: Crosslinking experiments suggest that SurA binds to uOmpA$_{171}$ with a delocalized interface.***

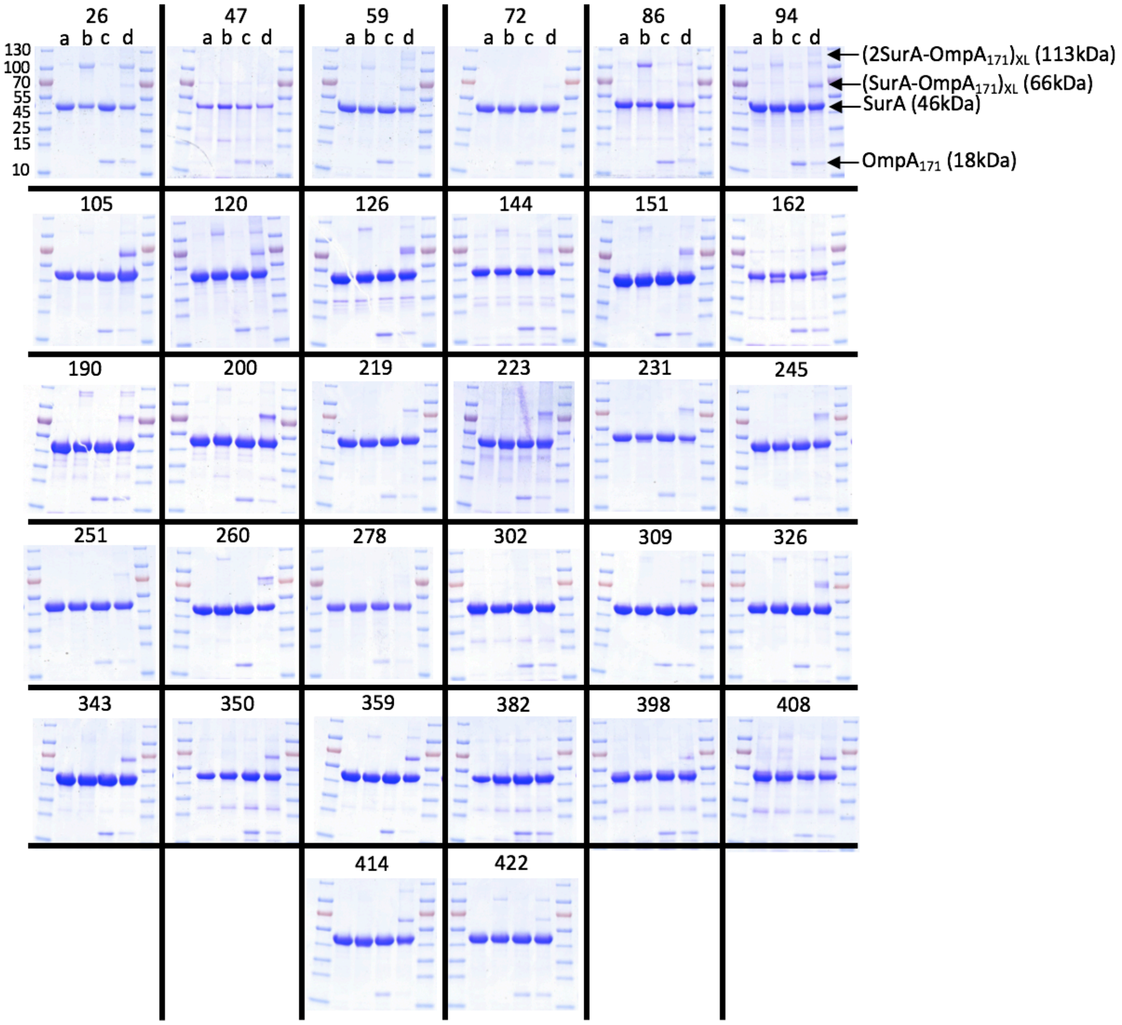

SurA$_{pAF}$ (25 μM) with or without uOmpA$_{171}$ (5 μM) was reconstituted in 20 mM Tris (pH 8), and exposed (or not) to UV light for 5 min. The resulting photo-products were then resolved and analyzed by SDS-PAGE. Representative SDS-PAGE analysis for crosslinking experiments between 36 SurA$_{pAF}$ variants and uOmpA$_{171}$ are shown. For each gel, the lanes are loaded as follows: SurA alone (**a** (-UV) and **b** (+UV)); SurA + uOmpA$_{171}$ mixture (**c** (-UV) and **d** (+UV)). Prior to UV exposure, the SurA variants and uOmpA$_{171}$ are observed as bands with apparent molecular weights of 46 kDa and 18 kDa, respectively. After UV exposure, some variants show a higher apparent molecular weight band (*i.e.*, "Complex"). The migration positions of the one-to-one and two-to-one species are shown for SurA$_{94,pAF}$.

#### *Figure S3. Crosslinking Efficiency of the Non-Cognate Client OmpLA is Low*

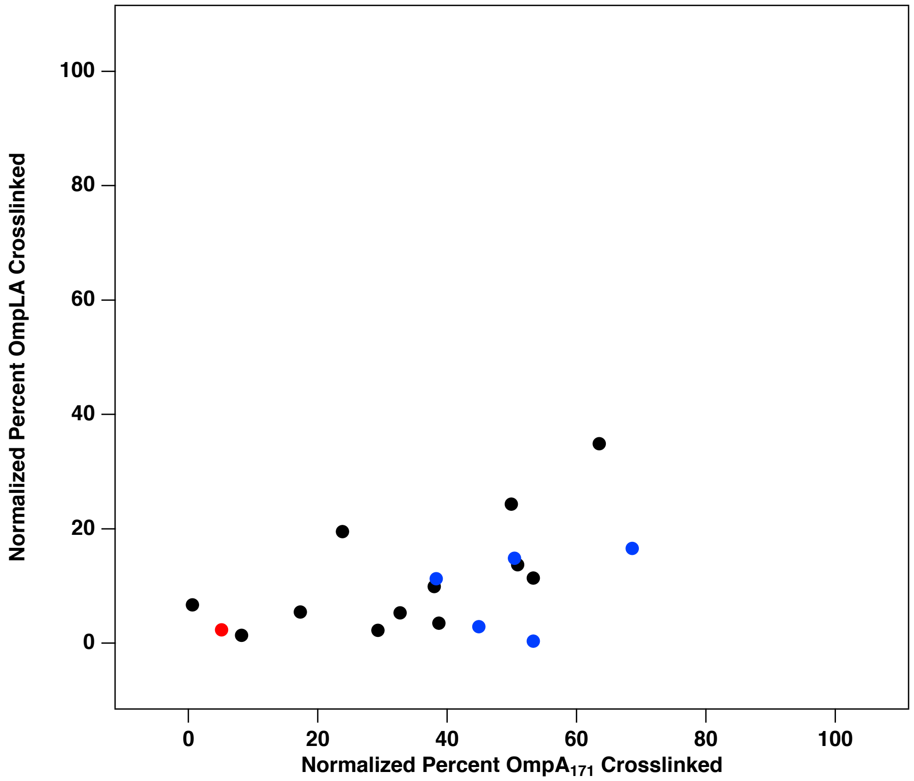

Normalized crosslinking efficiencies of the 36 $SurA_{pAF}$ variants to $uOmpA_{171}$ and uOmpLA. Each point represents an individual $SurA_{pAF}$ variant (colored by the domain of which it is part). The crosslinking efficiencies of SurA to uOmpLA were much lower and do not correlate to their efficiencies to $uOmpA_{171}$, consistent with a hypothesis that uOmpLA is not a substrate for SurA and crosslinks to it reflect non-specific associations.

***Figure S4: Compact apo SurA structures do not colocalize high-efficiency crosslinking sites***

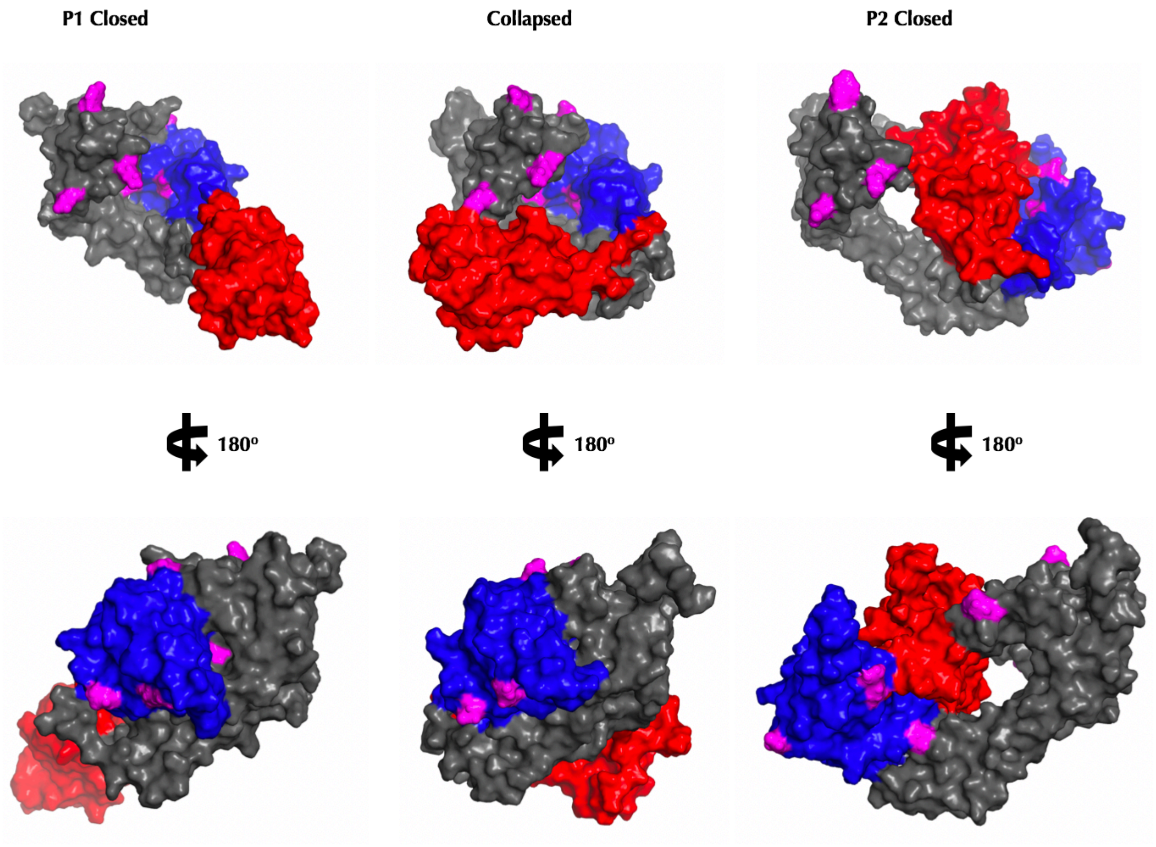

SurA is shown in a surface representation with domains in each model colored as in Figure 1A. The positions of the eight high-efficiency crosslinking sites are shown in magenta. In the P1 closed conformation (identified in x-ray crystallography), the pink residues found on the core and P1 domains are on opposite sides of the protein (shown by 180° rotation), which does not allow for a distinct uOMP interaction site to be identified. The collapsed conformation (where both P1 and P2 domains are bound to the core domain) and the P2-closed conformation (where P2 is bound to the core domain and P1 is structurally isolated) have both been recently shown to possibly exist in solution (11). Models created of these conformations also did not allow for a distinct uOMP interaction site to be identified.

***Figure S5. Structural Analysis of uOMP Binding Groove in "open" SurA***

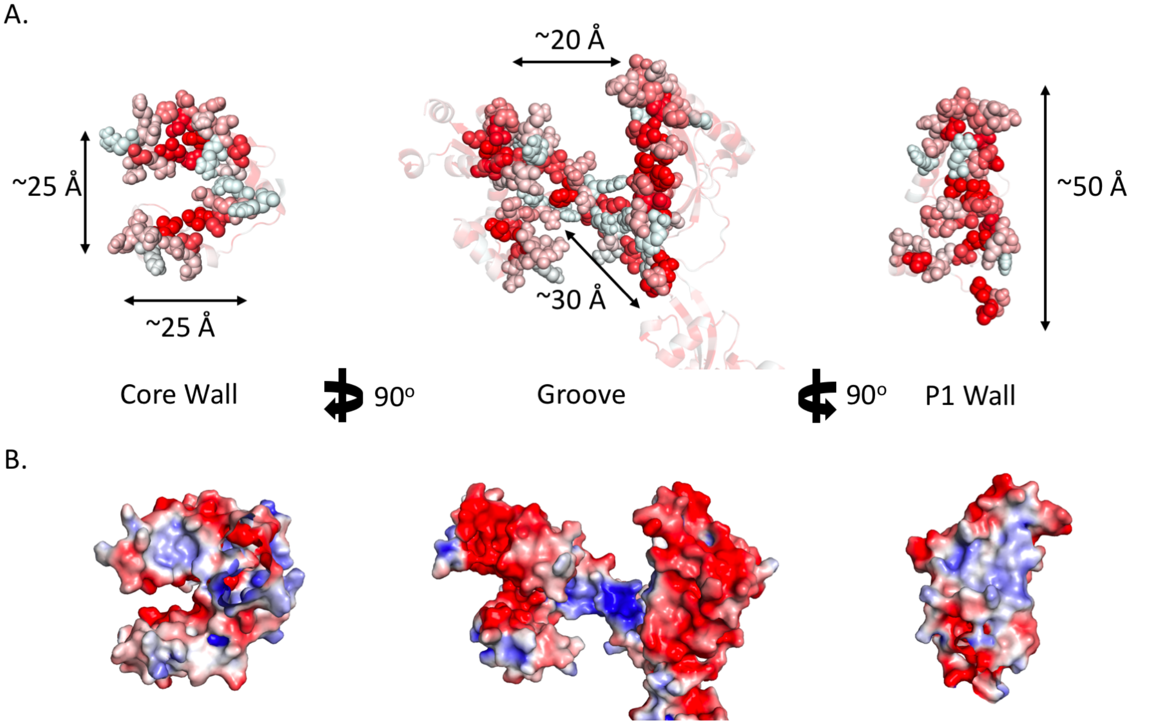

In the "open" conformation of SurA, a groove forms that contains hydrophobic patches and is electropositive in nature. In the middle of each panel, the groove is shown from a top-down perspective; to the left and right are 90° rotations, illustrating the contributions of the core domain and the P1 domain to create the "walls" of the groove. Panel **A** shows the groove in a space-filling representation, with residues colored based on hydrophobicity (red is most hydrophobic). The dimensions of the groove are also denoted in this panel, including the length of the floor of the groove, which is made of the C-terminal helix of the core domain. Panel **B** show the groove in a surface representation colored based on the electrostatic potential of the surface (± 3k/T). The floor and walls of the groove are positively charged (blue), while the surface near the top of the groove is negatively charged (red).

***Figure S6. Scattering Contribution for SurA and uOmpA$_{171}$ as a Function of %$D_2O$***

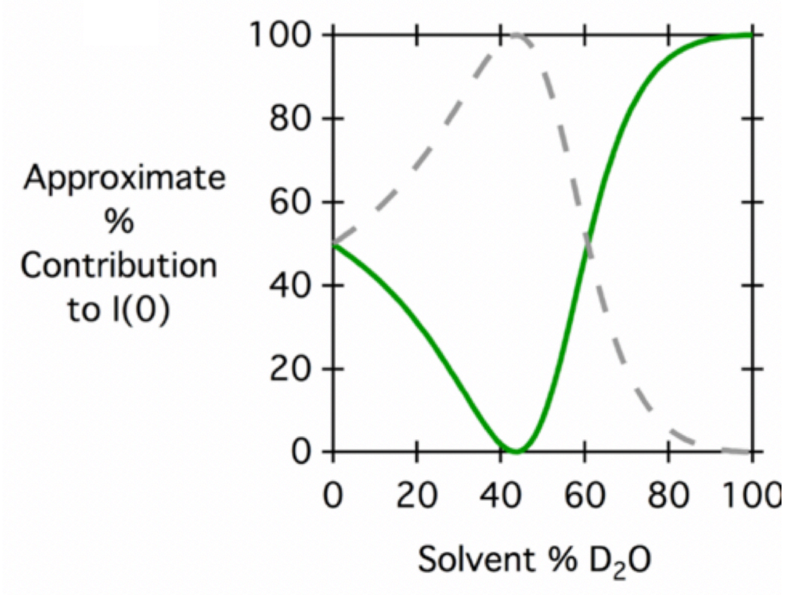

The contributions to the overall scattering intensity derived from protonated-SurA (green, solid line) and perdeuterated-uOmpA$_{171}$ (gray, dashed line) are plotted as a function of percent $D_2O$ in the sample buffer. In our SANS experiments, we utilize two conditions 0% $D_2O$ (where each protein contributes equally to scattering) and 30% $D_2O$ (where uOmpA$_{171}$ contributes 84% of the scattering intensity).

***Figure S7: Flowchart and Examples of apo uOmpA$_{171}$ structural models***

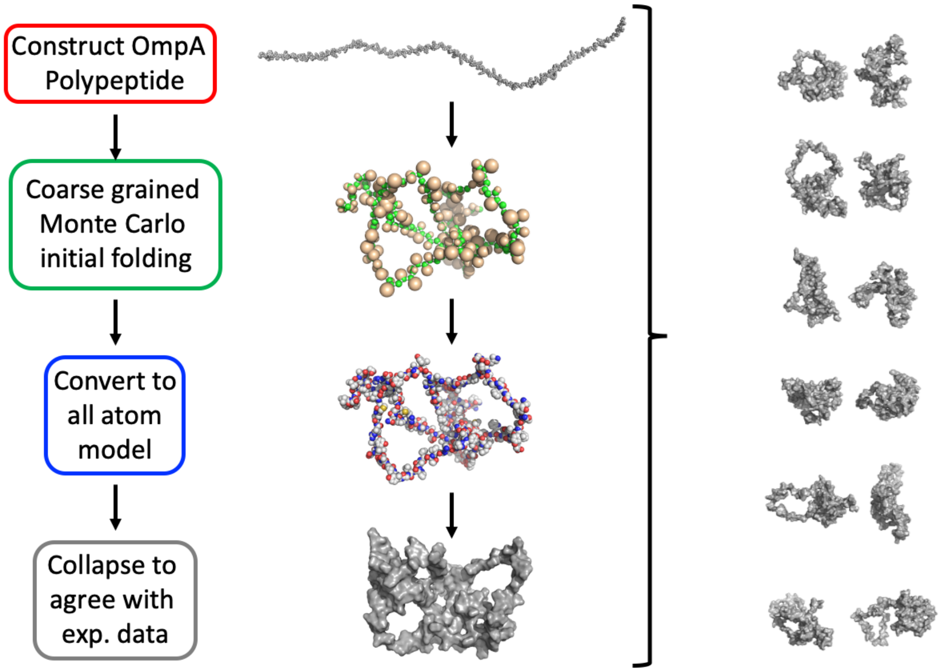

Twelve independent uOmpA$_{171}$ models (residues 22-192) were created using the following protocol and are shown in the panel on the right. An initial, extended OmpA polypeptide (φ=-78˚, ψ=149˚) was constructed where amino acid residues were converted to a coarse-grained model with single pseudo-atom side chains. Torsion angles were altered using a Monte Carlo approach to obtain a randomly folded, but relatively expanded model (green and gold spheres). For the Monte Carlo procedure, new phi/psi values for a randomly chosen residue were attempted. The structure was filtered for atomic overlap, and the Metropolis criterion was applied with a scoring function that included residue-specific Ramachandran propensities (12) and backbone-backbone hydrogen bonding (13). After 2000 moves the structures were considered sufficiently folded for further collapse with molecular dynamics *in vacuo*. The initially folded course-grained model was converted to an all atom model (red, blue, white spheres) and further collapsed using molecular dynamics simulations in generalized Born implicit solvent with the collective variables module in NAMD (grey molecular surface). For the molecular dynamics collapse, 200 steps of energy minimization in the CHARMM22 force field were followed by 50,000 to 150,000 steps of MD with implicit solvent alpha cutoff=12.0 Å, [ion]=0.3M, non-bonded cutoff=14.0, switching starting at 13.0 and 2 fs time step. Langevin dynamics was used with a damping coefficient of 1 for temperature control (NVT). A collective variables radius of gyration biasing potential (lower wall=20.0 Å, upper wall=25Å) was used for final collapse using collective variables to drive molecular dynamics simulations (14). HullRad was used to calculate radius of gyration and sedimentation coefficient during the simulation and a structure was saved when $R_G$ = 24.95±1.06 Å and s = 1.641± 0.078 (15).

***Figure S8: $R_G$ vs. s-value for intrinsic uOmpA$_{171}$ models***

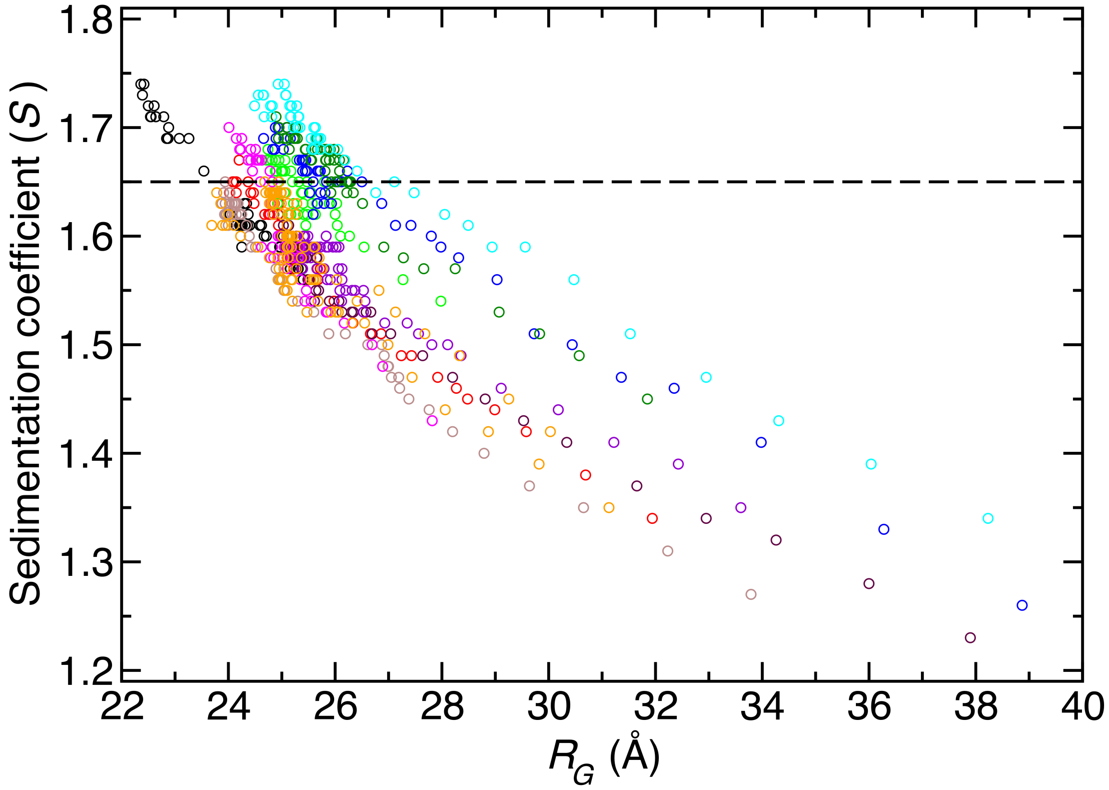

Each uOmpA model created using coarse-grained Monte Carlo folding was further collapsed using the collective variables module in NAMD under implicit solvent conditions. The relationship of calculated sedimentation coefficient and $R_G$ during collapse is plotted with different colored circles for each of the 12 initial models. Structures with $s \approx 1.65$ were chosen as representative structures of the intrinsic uOmpA$_{171}$ conformation in solution without denaturant or chaperones present.

*Figure S9: HADDOCK Docking of uOmpA$_{171}$ segments to SurA*

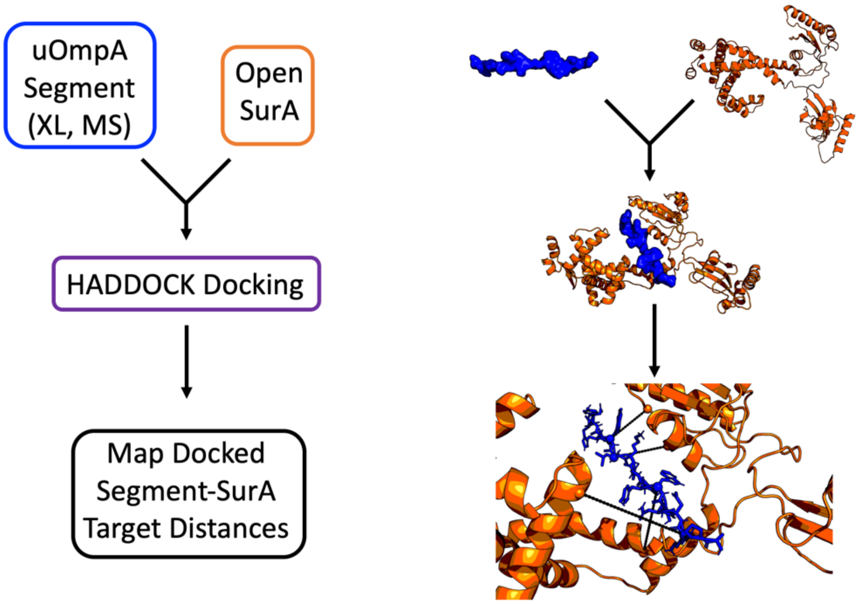

Four OmpA$_{171}$ amino acid segments (2-21, 54-73, 84-104, 115-132) which contain all of the SurA binding segments identified with XL-MS were modelled as extended polypeptides ($\varphi$=-78°, $\psi$=149°, blue spheres, top right panel). These segments were individually docked to the open form of SurA (orange ribbon) using HADDOCK(16). High ranking docked segment-SurA complexes were inspected to obtain target distances between adjacent uOmpA segment and SurA residues (bottom right panel). These target distances were used to dock full length uOmpA$_{171}$ models to "open" SurA (Figure S10).

***Figure S10: Docking uOmpA$_{171}$ to SurA and Expanding Bound uOmpA$_{171}$ Flowchart***

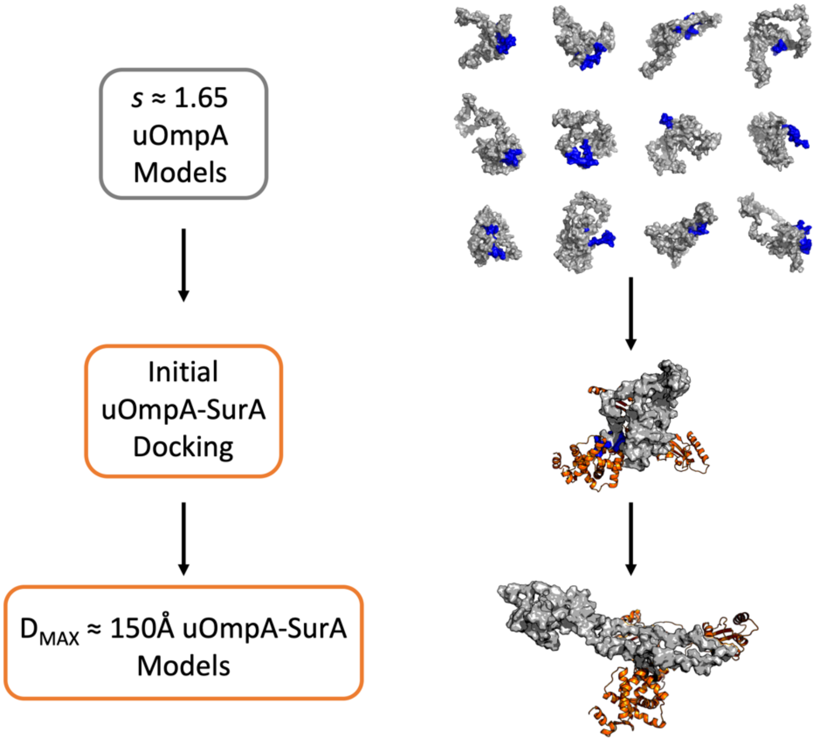

Individual uOmpA models with the corresponding crosslinking segment on the protein surface (blue, top right panel) were identified and docked to open form SurA using the target distances obtained from HADDOCK docking (middle panel). The collective variables module in NAMD with implicit solvent conditions was used to remove atomic clash during docking. Additional uOmpA-SurA models were made by expanding the maximum dimensions of docked uOmpA to ~150 Å as suggested by SANS analysis in 30% $D_2O$ (lower panel).

***Figure S11: Example structural models of SurA-uOmpA$_{171}$ complex***

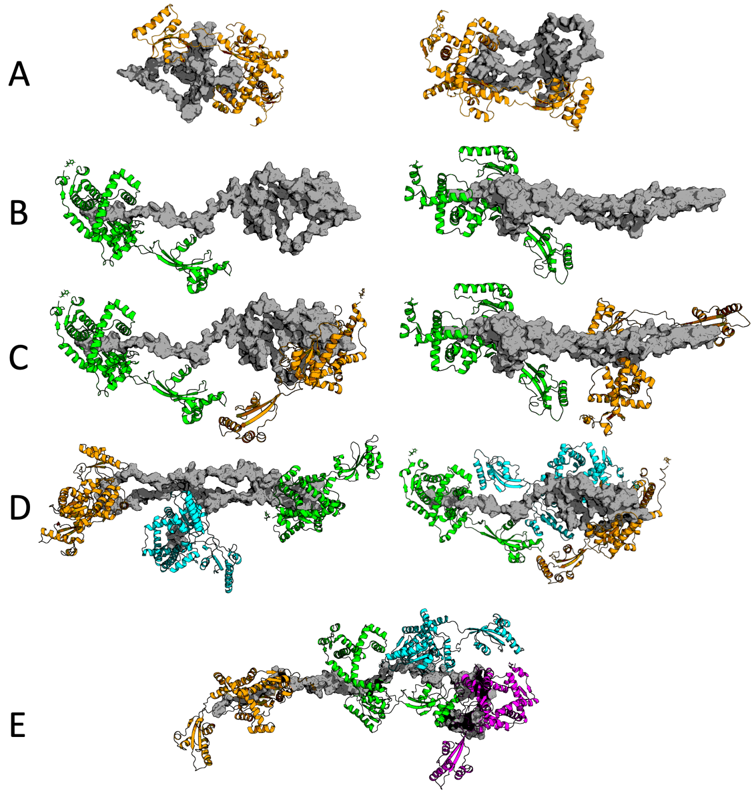

Representative snapshots of SurA-uOmpA$_{171}$ models used in the basis set for SANS analysis. **(A)** One SurA docked to non-expanded uOmpA$_{171}$ (showing 2 of 6 total); **(B)** one SurA docked to expanded uOmpA$_{171}$ (2 of 17); **(C)** two SurA docked to expanded uOmpA$_{171}$ (2 of 13); **(D)** three SurA docked to expanded uOmpA$_{171}$ (2 of 3); **(E)** four SurA docked to expanded uOmpA$_{171}$ (1 of 1).

*Figure S12: DSBU XL-MS Crosslinking shows client uOMPs bind in the SurA groove*

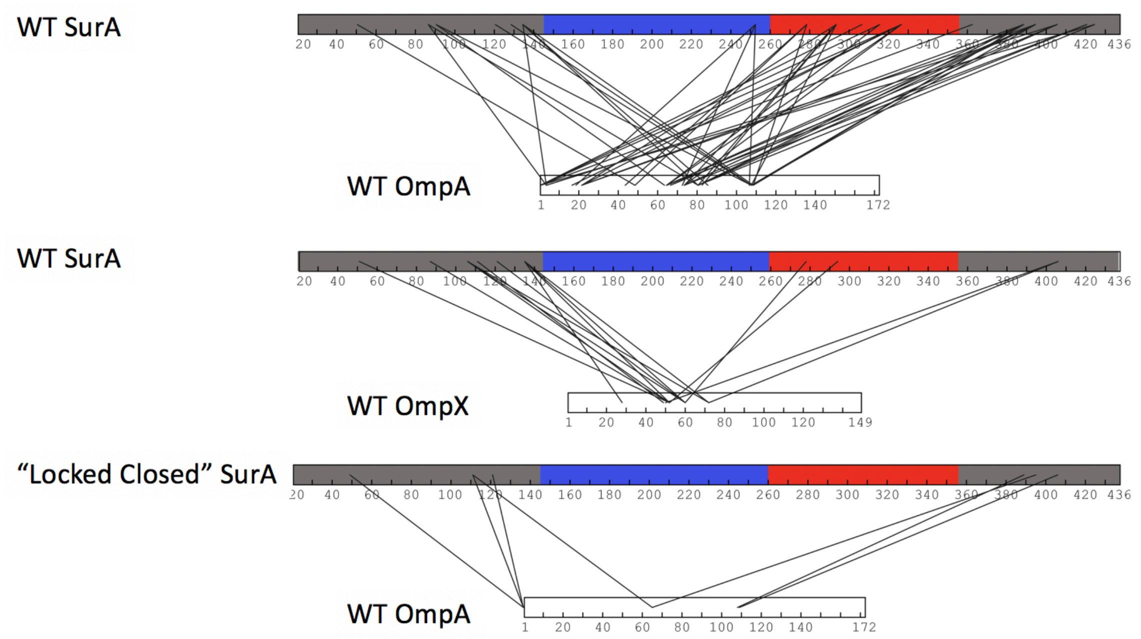

DSBU crosslinks found between SurA and client uOMPs are shown (17). Sequences are shown as bars, with the SurA sequences colored based on the domain architecture outlined in Figure 1A. Lines between sequences represent a DSBU crosslink between residues, as determined by XL-MS (see Supplementary Data 3-5). We find that many residues on SurA or uOmpA$_{171}$ crosslink to multiple residues on the other protein, which is denoted by many lines originating from a single point of origin on the sequence diagrams above. The top two diagrams represent WT SurA crosslinking to each client uOMPs, uOmpA$_{171}$ and uOmpX. The bottom diagram contains the "Locked-Closed" SurA variant (P61C/A218C) in which a disulfide bond between the core and P1 domains inhibits the uOMP binding groove from forming. The difference in the total amount of crosslinks found comparing WT SurA and "Locked Closed" SurA mixed with uOmpA$_{171}$ shows that the formation of the groove is essential for efficient client uOMP binding.

***Figure S13. Clusters among DSBU crosslinks and three SurA•uOmpA$_{171}$ binding modes***

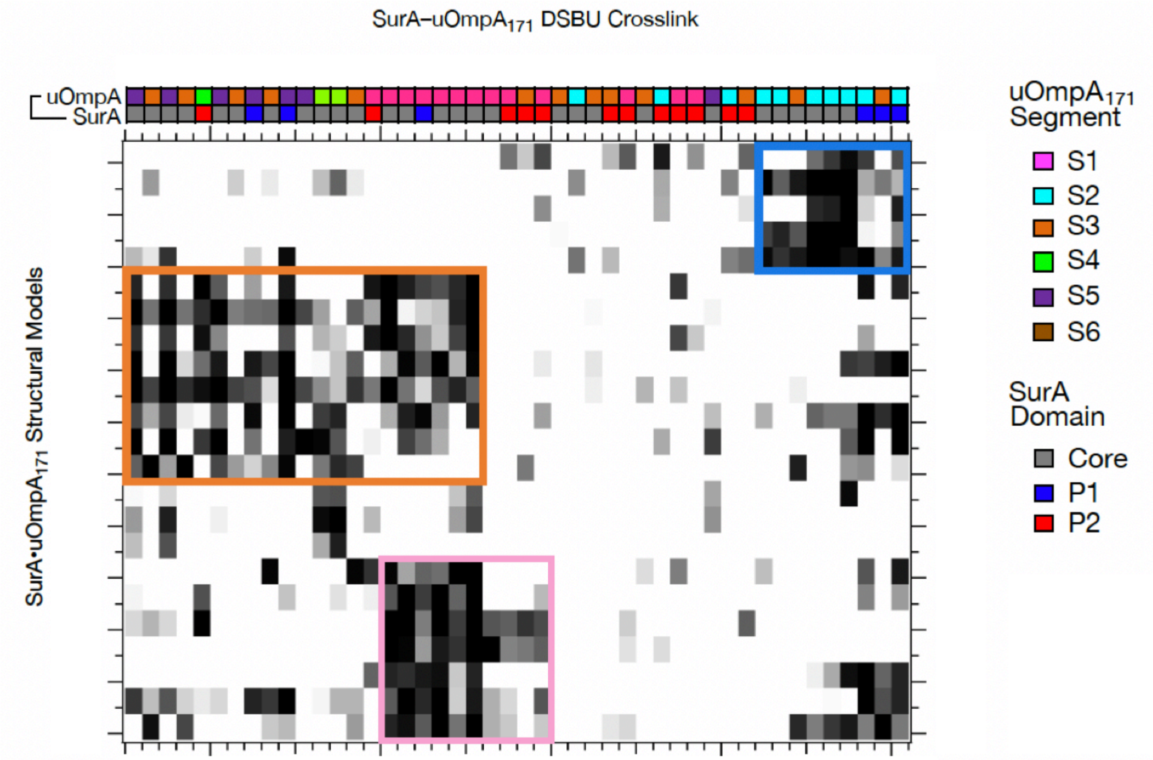

Spectral biclustering reveals a natural grouping among structural models (rows) that are collectively consistent with subsets of crosslinks (columns) (18). Structures divide into a pink cluster (wherein SurA binds segment 1; 7 models), a blue cluster (wherein SurA binds segment 2; 5 models), and an orange cluster (wherein SurA binds uOmpA at segment 1 and either segments 3–5; 8 models). A few of the structures are not well explained by any of the crosslinks. Crosslinks divide into a pink cluster (which support the pink structures, and to a lesser extent, the orange structures), a blue cluster (which support the blue structures), and an orange cluster (which support the orange structures). The blue cluster has a sub-cluster (blue-pink) which is also consistent with some of the pink structures. Twelve of the crosslinks are not well explained by any of the 23 SurA•uOmpA$_{171}$ models. Crosslinks are annotated with colors, with the SurA site represented by the domain it is on and the uOmpA$_{171}$ site represented by the nearest binding segment. The grayscale of the matrix represents the scores for a given crosslink in a given structure (white = 0, black = 100; see SI methods for explanation); all SASDs and scores are in Supplementary Data 6.

***Figure S14. SANS Profile of SurA$_{105,pAF}$-uOmpA$_{171}$ crosslinked complex in 0% $D_2O$***

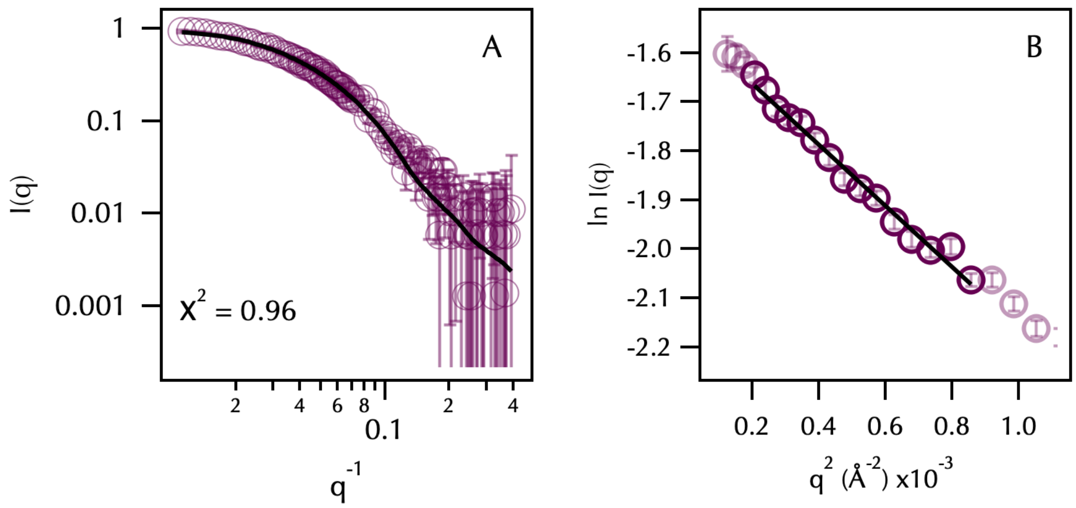

(**A**) SANS data for (SurA$_{105,pAF}$–uOmpA$_{171}$)$_{XL}$ with SurA protonated and uOmpA$_{171}$ perdeuterated, for the 0% $D_2O$ buffer condition. Error bars represent the standard error of the mean with respect to the number of pixels used in the data averaging. The black lines through the data are the average waves from triplet basis setting with the $\chi^2$ for the fits. (**B**) Guinier regions of these data with a linear fit (values for the Guinier fits Table S3).

***Figure S15. SDS PAGE of SEC Fractions indicates crosslinked SurA-uOmpA$_{171}$ cannot be fully separated from excess SurA.***

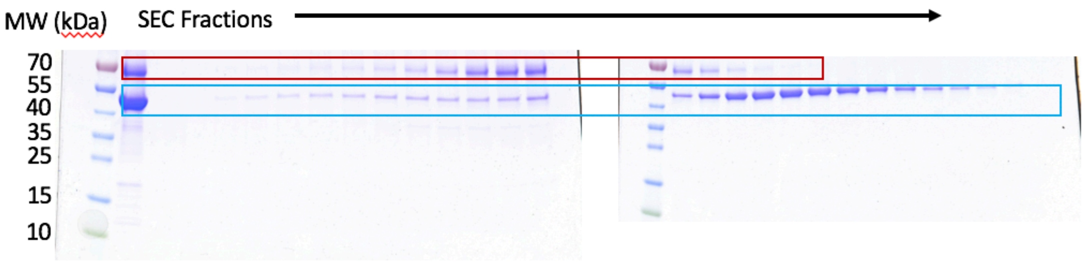

SDS PAGE gel shows the crosslinked SurA$_{105,pAF}$-uOmpA171 sample that was injected on to the SEC column (first lane on left gel). Further lanes are 0.5mL fractions collected from the SEC run, with the crosslinked complex (red box) and free SurA (blue box) highlighted. Because of the small change in size (45 vs 65 kDa), and the possibility that free SurA is interacting with the crosslinked complex, we could not completely separate the two species using SEC.

***Figure S16. NMR Determination of Deuteration Level of uOmpA used in some SANS Experiments.***

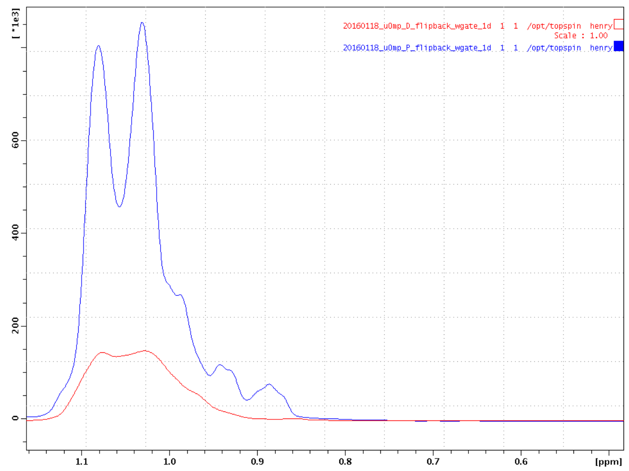

OMP deuteration was estimated to be 80%. We collected 1D $^{1}H$ spectra on both protonated and deuterated uOmp$A_{171}$ (50 mM) in 8 M Urea, 20 mM Tris, 10% $D_2O$ at 35 °C, on a 600 MHz Bruker Avance II spectrometer. Water suppression was achieved using a flipback-watergate sequence and a buffer purging pulse was included to minimize the large urea peak. Each spectrum was collected using 128 scans, a recycle delay of 1.5 s, and acquisition time of 60 ms/FID. Data was processed and analyzed using TopSpin 2.1. The spectra were aligned using the amide resonance peaks, and the baseline of the methyl peaks was corrected using a $5^{th}$ order polynomial. After baseline correction, the methyl peak volumes were integrated using TopSpin, and the integrated intensities of the methyl peaks in both the protonated and deuterated samples were compared. The loss of intensity in the methyl peaks between the two samples was used to estimate the deuteration level of deuterated-uOmp$A_{171}$.

### Supplemental Tables

#### *Table S1. SurA$_{pAF}$ Variant Crosslinking Efficiencies*

| | uOmpA (n=3-5) | uOmpX (n=3) | uOmpLA (n=2) |
|---|---|---|---|
| pAF Residue | Percent Crosslinked | Percent Crosslinked | Percent Crosslinked |
| D26 | 23.8 ± 6.3 | 23.7 ± 6.9 | 19.5 ± 3.4 |
| Q47 | 0.6 ± 6.0 | 17.3 ± 11.4 | 6.7 ± 3.5 |
| Q59 | 63.5 ± 7.0 | 62.1 ± 4.8 | 34.9 ± 7.5 |
| E72 | 32.7 ± 5.6 | 28.7 ± 9.0 | 5.3 ± 6.4 |
| K86 | 38.7 ± 9.8 | 24.7 ± 7.2 | 3.5 ± 6.9 |
| E94 | 49.9 ± 9.5 | 41.6 ± 8.2 | 24.3 ± 6.4 |
| K105 | 53.3 ± 13.1 | 51.8 ± 11.4 | 11.4 ± 11.1 |
| Y120 | 50.9 ± 8.7 | 43.3 ± 4.1 | 13.7 ± 6.1 |
| N126 | 17.3 ± 12.9 | 18.2 ± 13.3 | 5.5 ± 3.6 |
| N144 | 38.0 ± 17.8 | 31.7 ± 8.4 | 9.9 ± 3.7 |
| T151 | 29.3 ± 9.9 | 29.6 ± 2.7 | 2.3 ± 5.2 |
| Q162 | 11.9 ± 6.2 | 10.6 ± 4.6 | n/a |
| D190 | 20.9 ± 8.7 | 21.6 ± 5.3 | n/a |
| R200 | 44.9 ± 12.0 | 33.9 ± 9.6 | 4.1 ± 6.1 |
| H219 | 38.3 ± 8.3 | 33.9 ± 4.1 | 2.9 ± 4.0 |
| Q223 | 50.4 ± 9.9 | 47.9 ± 3.2 | 11.3 ± 6.0 |
| M231 | 53.3 ± 14.2 | 54.5 ± 4.4 | 14.9 ± 8.7 |
| Q245 | 68.6 ± 6.6 | 68.9 ± 5.7 | 16.6 ± 7.3 |
| K251 | 21.6 ± 4.7 | 18.2 ± 4.3 | n/a |
| R260 | 67.7 ± 10.3 | 69.6 ± 5.1 | n/a |
| K278 | 18.7 ± 12.1 | 22.9 ± 5.7 | n/a |
| Q302 | 8.4 ± 8.2 | 3.3 ± 4.6 | n/a |
| Q309 | 12.3 ± 11.7 | 7.2 ± 3.4 | n/a |
| K326 | 41.1 ± 13.0 | 29.7 ± 4.9 | n/a |
| W343 | 33.1 ± 5.9 | 19.1 ± 5.3 | n/a |
| D350 | 25.9 ± 11.5 | 23.9 ± 9.9 | n/a |
| R359 | 33.9 ± 13.2 | 33.6 ± 7.2 | n/a |
| D382 | 5.1 ± 13.8 | 11.3 ± 5.0 | 2.3 ± 3.5 |
| Y398 | 10.8 ± 13.5 | 11.4 ± 7.9 | n/a |
| E408 | 31.3 ± 4.7 | 16.9 ± 5.1 | n/a |
| M414 | 62.6 ± 8.5 | 31.1v± 15.1 | n/a |
| Y422 | 8.2 ± 9.8 | 14.2 ± 3.6 | 1.39 ± 6.0 |

Crosslinking efficiencies for SurApAF variants crosslinked to three different outer membrane proteins. The errors reported for uOmpA$_{171}$ and uOmpX are standard deviation; the errors reported for the OmpLA values is the standard error of the mean. Values were considered not applicable if values were indistinguishable from controls lacking SurA. Values were corrected for total intensity of uOmpA$_{171}$ lost due to UV alone (uOmpA$_{171}$ = 21%, uOmpX = 13%, uOmpLA = 22%).

*Table S2. Description of apo SurA Conformational Variants*

| | |
|---|---|
| Open | This conformation has both P1 and P2 domains open away from the core domain. It was built from 1M5Y, and the relative orientation of P1 to the core is the same as in 2PV3. |
| P1 closed | This conformation is based on the 1M5Y crystal structure. Missing loops and a C-terminal His tag were added using Modeller.(19) |
| P2 closed | The core-P1 orientation is based on the 2PV3 dimer structure, and the P2 is collapsed in the binding groove between core and P1. |
| Collapsed | The P1-core domain structures are the same as 1M5Y, with the P2 domain collapsed onto the core. |

***Table S3. Parameters from Guinier Analysis of SANS Data***

| | **Concentration (mg mL$^{-1}$)** | **I(0) (cm$^{-1}$)** | **$R_G$ (Å)** | **$R_G$*q Range** |
|---|---|---|---|---|
| ($SurA_{105}$-$uOmpA_{171}$)$_{XL}$ (in 0% $D_2O$ Hydrogenated SurA Deuterated $uOmpA_{171}$) | 3.0 | 0.217 ± 0.002 | 44.0 ± 0.7 | 0.585 – 1.239 |
| | | 0.215 ± 0.002 | 43.2 ± 0.7 | 0.621 – 1.265 |
| | | 0.212 ± 0.003 | 42.6 ± 0.9 | 0.747 – 1.245 |
| ($SurA_{105}$-$uOmpA_{171}$)$_{XL}$ (in 30% $D_2O$ Hydrogenated SurA Deuterated $uOmpA_{171}$) | 3.0 | 0.054 ± 0.002 | 46 ± 3 | 0.615 – 1.254 |
| | | 0.053 ± 0.002 | 45 ± 3 | 0.642 – 1.213 |
| | | 0.053 ± 0.002 | 45 ± 3 | 0.696 – 1.271 |

Summary of the fitting parameters derived from Guinier fitting using a range of $qR_G$ values for analysis. For $I(0)$ and $R_G$ values, errors indicate the standard deviations from fitting. Rows shown in grey are included in Table S4.

***Table S4. Parameters from P(r) Analysis of SANS Data***

| | Concentration (mg mL$^{-1}$) | $D_{max}$ (Å) | I(0) (cm$^{-1}$) | $R_G$ (Å) | q Range (Å$^{-1}$) |
|---|---|---|---|---|---|
| Prot-SurA$_{pAF}^{105}$ crosslinked to deut-uOmpA$_{171}$ (0% $D_2O$) | 3.0 | 140 | 0.209 ± 0.001 | 42.0 ± 0.3 | 0.01436 – 0.1977 |
| | | 150 | 0.212 ± 0.002 | 43.8 ± 0.5 | 0.01436 – 0.1977 |
| | | 160 | 0.215 ± 0.002 | 45.1 ± 0.6 | 0.01436 – 0.1977 |
| Prot-SurA$_{pAF}^{105}$ crosslinked to deut-uOmpA$_{171}$ (30% $D_2O$) | 3.0 | 140 | 0.051 ± 0.001 | 44 ± 1 | 0.01436 – 0.1496 |
| | | 150 | 0.052 ± 0.001 | 45 ± 1 | 0.01436 – 0.1496 |
| | | 160 | 0.053 ± 0.001 | 46 ± 2 | 0.01436 – 0.1496 |
| | | 170 | 0.054 ± 0.001 | 47 ± 2 | 0.01436 – 0.1496 |

Fitting parameters derived from generation of distance distribution functions, P(r) vs. r curves, for all SANS datasets using the specified $D_{Max}$ values. For I(0) and $R_G$ values, errors indicate the standard deviations from fitting. Rows shown in grey are included in Table S3.

***Table S5. Summary of all XL-MS and pXL-MS injections and FDR cut-offs.***

| | **SurA Variant** [a] | **uOMP Variant** [b] | **XL**[c] | **Digest** [d] | ***n*** [e] | **$N_{PSM}$ inter** [f] | **inter FDR cutoff**[g] | **$N_{PSM}$ Intra**[h] | **intra FDR cutoff**[i] |
|---|---|---|---|---|---|---|---|---|---|
| 1 | SurApaf105 | OmpA171 | paf | Trypsin | 1 | 9 | 14.6 | 15 | 17 |
| 2 | SurApaf105 | OmpA171 | paf | Glu-C | 1 | 16 | 46.7 | 37 | 34.9 |
| 3 | SurApaf105 | OmpA171 | paf | Glu-C | 2 | 21 | 45.8 | 13 | 32.3 |
| 4 | surApaf245 | OmpA171 | paf | Trypsin | 1 | 7 | 15.3 | 2 | 14.5 |
| 5 | surApaf245 | OmpA171 | paf | Trypsin | 2 | 9 | 15 | 2 | 17 |
| 6 | surApaf245 | OmpA171 | paf | Glu-C | 1 | 23 | 26.2 | 1 | 34.4 |
| 7 | surApaf245 | OmpA171 | paf | Glu-C | 2 | 18 | 22.1 | 2 | 40.4 |
| 8 | surApaf260 | OmpA171 | paf | Trypsin | 1 | 17 | 17 | 1 | 14 |
| 9 | surApaf260 | OmpA171 | paf | Trypsin | 2 | 14 | 16.2 | 2 | 12 |
| 10 | surApaf260 | OmpA171 | paf | Glu-C | 1 | 7 | 14.5 | 1 | 19.8 |
| 11 | surApaf260 | OmpA171 | paf | Glu-C | 2 | 7 | 15.6 | 1 | 14.9 |
| 12 | surApaf26 | N/a | paf | Trypsin | 1 | N/a | N/a | 0 | 11.2 |
| 13 | surApaf26 | N/a | paf | Trypsin | 2 | N/a | N/a | 0 | 1.1 |
| 14 | surApaf26 | N/a | paf | Glu-C | 1 | N/a | N/a | 0 | 11 |
| 15 | surApaf26 | N/a | paf | Glu-C | 2 | N/a | N/a | 0 | 12.3 |
| 16 | surApaf26 | OmpA171 | paf | Trypsin | 1 | 0 | N/a | 0 | N/a |
| 17 | surApaf26 | OmpA171 | paf | Trypsin | 2 | 0 | N/a | 0 | 0.9 |
| 18 | surApaf26 | OmpA171 | paf | Glu-C | 1 | 1 | 14.9 | 0 | 12.6 |
| 19 | surApaf26 | OmpA171 | paf | Glu-C | 2 | 0 | 0.8 | 0 | 5.3 |
| 20 | surApaf59 | OmpA171 | paf | Trypsin | 1 | 49 | 23.1 | 25 | 28.8 |
| 21 | surApaf59 | OmpA171 | paf | Trypsin | 2 | 37 | 33.6 | 14 | 27.5 |
| 22 | surApaf59 | OmpA171 | paf | Glu-C | 1 | 44 | 43.6 | 6 | 41.4 |
| 23 | surApaf59 | OmpA171 | paf | Glu-C | 2 | 36 | 36.8 | 3 | 25.3 |
| 24 | surApaf94 | OmpA171 | paf | Trypsin | 1 | 24 | 23.7 | 8 | 18 |
| 25 | surApaf94 | OmpA171 | paf | Trypsin | 2 | 22 | 18.8 | 6 | 12.7 |
| 26 | surApaf94 | OmpA171 | paf | Glu-C | 1 | 17 | 13.8 | 2 | 16.6 |
| 27 | surApaf94 | OmpA171 | paf | Glu-C | 2 | 14 | 17.2 | 2 | 16.8 |
| 28 | surApaf120 | OmpA171 | paf | Trypsin | 1 | 24 | 18.7 | 14 | 40.8 |
| 29 | surApaf120 | OmpA171 | paf | Trypsin | 2 | 23 | 26.5 | 15 | 17.9 |
| 30 | surApaf120 | OmpA171 | paf | Glu-C | 1 | 55 | 23.5 | 8 | 23.4 |
| 31 | surApaf120 | OmpA171 | paf | Glu-C | 2 | 25 | 19.6 | 5 | 23.8 |
| 32 | surApaf223 | OmpA171 | paf | Trypsin | 1 | 2 | 12.7 | 17 | 16.9 |
| 33 | surApaf223 | OmpA171 | paf | Trypsin | 2 | 2 | 10 | 10 | 16.6 |
| 34 | surApaf223 | OmpA171 | paf | Glu-C | 1 | 3 | 20.1 | 29 | 20.3 |
| 35 | surApaf223 | OmpA171 | paf | Glu-C | 2 | 5 | 20.7 | 15 | 25.7 |
| 36 | surApaf231 | OmpA171 | paf | Trypsin | 1 | 6 | 14 | 2 | 11.7 |
| 37 | surApaf231 | OmpA171 | paf | Glu-C | 1 | 19 | 13.3 | 2 | 16.7 |
| 38 | surApaf422 | OmpA171 | paf | Trypsin | 1 | 0 | 15.4 | 1 | 18.4 |
| 39 | surApaf422 | OmpA171 | paf | Trypsin | 2 | 0 | 10.7 | 0 | 18.7 |
| 40 | surApaf422 | OmpA171 | paf | Glu-C | 1 | 0 | 12.9 | 0 | 22.1 |
| 41 | surApaf422 | OmpA171 | paf | Glu-C | 2 | 1 | 10.3 | 2 | 34.5 |
| 42 | SurApaf105 | OmpX | paf | Trypsin | 1 | 11 | 10.9 | 0 | 11 |
| 43 | SurApaf105 | OmpX | paf | Glu-C | 1 | 6 | 13.7 | 3 | 18.1 |
| 44 | SurApaf245 | OmpX | paf | Trypsin | 1 | 3 | 4.5 | 0 | 2.4 |
| 45 | SurApaf245 | OmpX | paf | Glu-C | 1 | 7 | 5.2 | 0 | 14.7 |

| | | | | | | | | | |
|---|---|---|---|---|---|---|---|---|---|
| 46 | SurApaf59 | OmpX | paf | Trypsin | 1 | 10 | 8.3 | 3 | 10.5 |
| 47 | SurApaf59 | OmpX | paf | Glu-C | 1 | 5 | 11.8 | 1 | 26.9 |
| 48 | SurApaf260 | OmpX | paf | Trypsin | 1 | 3 | 8.1 | 0 | 0.1 |
| 49 | SurApaf260 | OmpX | paf | Glu-C | 1 | 4 | 10.6 | 2 | 19.9 |
| 50 | SurApaf26 | OmpX | paf | Trypsin | 1 | 0 | 1.7 | 0 | 13.5 |
| 51 | SurApaf26 | OmpX | paf | Glu-C | 1 | 0 | 6 | 0 | 7.2 |
| 52 | SurApaf94 | OmpX | paf | Trypsin | 1 | 5 | 11.5 | 1 | 13.5 |
| 53 | SurApaf94 | OmpX | paf | Glu-C | 1 | 3 | 12.6 | 1 | 29.1 |
| 54 | SurApaf120 | OmpX | paf | Trypsin | 1 | 16 | 15.2 | 18 | 22.8 |
| 55 | SurApaf120 | OmpX | paf | Glu-C | 1 | 0 | 7.8 | 0 | 13 |
| 56 | SurApaf223 | OmpX | paf | Trypsin | 1 | 6 | 4.5 | 15 | 15.1 |
| 57 | SurApaf223 | OmpX | paf | Glu-C | 1 | 6 | 12.9 | 10 | 34.2 |
| 58 | SurApaf231 | OmpX | paf | Trypsin | 1 | 7 | 10.3 | 6 | 12.7 |
| 59 | SurApaf231 | OmpX | paf | Glu-C | 1 | 5 | 8.9 | 1 | 13.5 |
| 60 | SurApaf422 | OmpX | paf | Trypsin | 1 | 0 | 0 | 0 | 9.9 |
| 61 | SurApaf422 | OmpX | paf | Trypsin | 2 | 0 | 0 | 0 | 0.1 |
| 62 | SurApaf422 | OmpX | paf | Glu-C | 1 | 0 | 9.2 | 0 | 9.3 |
| 63 | SurApaf422 | OmpX | paf | Glu-C | 2 | 0 | 8.7 | 0 | 16.3 |
| 64 | SurA WT | OmpA171 | DSBU | Trypsin | 1 | 10 | 0.2 | 58 | 2.2 |
| 65 | SurA WT | OmpA171 | DSBU | Trypsin | 2 | 16 | 1.1 | 34 | 1.9 |
| 66 | SurA WT | OmpA171 | DSBU | Glu-C | 1 | 24 | 0.9 | 25 | 1.2 |
| 67 | SurA WT | OmpA171 | DSBU | Glu-C | 2 | 28 | 1.5 | 37 | 2.3 |
| 68 | SurA WT | OmpX | DSBU | Trypsin | 1 | 4 | 0.5 | 24 | 7.3 |
| 69 | SurA WT | OmpX | DSBU | Trypsin | 2 | 7 | 0.5 | 35 | 0.6 |
| 70 | SurA WT | OmpX | DSBU | Glu-C | 1 | 6 | 0.7 | 9 | 4.1 |
| 71 | SurA WT | OmpX | DSBU | Glu-C | 2 | 3 | 0.4 | 5 | 0 |
| 72 | SurA PC2 | OmpA171 | DSBU | Trypsin | 1 | 0 | 0 | 29 | 1.7 |
| 73 | SurA PC2 | OmpA171 | DSBU | Trypsin | 2 | 1 | 0 | 23 | 0.3 |
| 74 | SurA PC2 | OmpA171 | DSBU | Glu-C | 1 | 4 | 0 | 11 | 1.5 |
| 75 | SurA PC2 | OmpA171 | DSBU | Glu-C | 2 | 3 | 1.8 | 10 | 1.2 |

[a]Variant of SurA indicating the position of *p*AF. PC2 designates the locked-closed double-mutant P61C/A218C. [b]Variant of unfolded Outer Membrane Protein. [c]Crosslinker used (paf = para-azidophenylalanine; DSBU = disuccinimidyl dibutyric urea). [d]Designates whether trypsin digest was conducted alone, or trypsin and Glu-C digests were conducted in serial. [e]Designates which replicate (for conditions done in technical duplicate). [f]Number of interprotein peptide-spectrum matches in this injection. [g]The score above which interprotein peptide-spectrum matches achieve an FDR < 0.01. [h]Number of intraprotein peptide-spectrum matches in this injection. [i]The score above which intraprotein peptide-spectrum matches achieve an FDR < 0.01.

***Table S6. Hydrodynamic Description of SurA-uOmpA$_{171}$ Complex Models***

| Model ID | Total $R_G$ | Total $D_{MAX}$ | uOmpA$_{171}$ $R_G$ | uOmpA$_{171}$ $D_{MAX}$ |
|---|---|---|---|---|
| o1s001 | 35.41 | 112.79 | 26.23 | 83.62 |
| o1s002 | 34.52 | 107.21 | 25.91 | 85.88 |
| o1s003 | 38.68 | 137.14 | 33.87 | 111.55 |
| o1s004 | 41.43 | 173.51 | 37.62 | 144.72 |
| o1s005 | 33.29 | 109.93 | 21.61 | 65.54 |
| o1s006 | 35.06 | 132.71 | 26.80 | 101.26 |
| o1s007 | 38.97 | 167.94 | 39.84 | 155.36 |
| o1s008 | 39.43 | 142.56 | 37.89 | 121.53 |
| o1s009 | 41.86 | 169.14 | 40.48 | 148.80 |
| o1s010 | 36.48 | 160.26 | 32.71 | 153.34 |
| o1s011 | 35.89 | 155.36 | 39.84 | 155.36 |
| o1s012 | 40.65 | 166.69 | 37.89 | 121.53 |
| o1s013 | 41.51 | 175.68 | 40.48 | 148.80 |
| o1s014 | 39.61 | 153.34 | 32.71 | 153.34 |
| o1s015 | 38.07 | 170.90 | 34.95 | 150.85 |
| o1s016 | 46.81 | 166.42 | 34.95 | 150.85 |
| o1s017 | 37.36 | 118.28 | 27.98 | 84.00 |
| o1s018 | 39.00 | 152.92 | 31.78 | 110.55 |
| o1s019 | 39.87 | 147.24 | 31.78 | 110.55 |
| o1s020 | 36.60 | 120.18 | 31.78 | 110.55 |
| o1s021 | 48.48 | 194.40 | 41.99 | 147.32 |
| o1s022 | 50.29 | 198.01 | 41.99 | 147.32 |
| o1s023 | 39.83 | 147.32 | 41.99 | 147.32 |
| o2s001 | 41.38 | 167.94 | 39.84 | 155.36 |
| o2s002 | 51.59 | 186.91 | 37.89 | 121.53 |
| o2s003 | 54.44 | 195.87 | 40.48 | 148.80 |
| o2s004 | 52.05 | 162.63 | 33.11 | 152.57 |
| o2s005 | 51.67 | 160.26 | 32.71 | 153.34 |
| o2s006 | 58.59 | 187.61 | 34.95 | 150.85 |
| o2s007 | 56.68 | 185.80 | 28.92 | 107.61 |
| o2s008 | 42.48 | 170.60 | 35.10 | 150.65 |
| o2s009 | 48.22 | 166.94 | 35.10 | 150.65 |
| o2s010 | 41.30 | 152.92 | 31.78 | 110.55 |
| o2s011 | 46.86 | 156.04 | 31.78 | 110.55 |
| o2s012 | 78.34 | 246.64 | 41.99 | 147.32 |
| o2s013 | 42.24 | 140.70 | 27.75 | 113.60 |
| o3s001 | 54.20 | 185.34 | 35.10 | 150.65 |
| o3s002 | 53.06 | 188.10 | 31.78 | 110.55 |
| o3s003 | 69.95 | 246.64 | 41.99 | 147.32 |
| o4s001 | 72.84 | 256.28 | 60.35 | 218.60 |

Highlighted models are found in the final ensemble of structures validated by XL-MS and 0% $D_2O$ SANS experiments.

***Table S7. Members of each Triplet of Structures that Fit the 0% $D_2O$ SANS Dataset.***

| Model 1 | Model 2 | Model 3 |
|---|---|---|
| "P1 closed" SurA (42.4) | o1s005 (14.2) | o1s016 (43.4) |
| "P1 closed" SurA (45.6) | o1s013 (35.1) | o1s016 (19.3) |
| "collapsed" SurA (40.2) | o1s001 (13.4) | o1s016 (46.4) |
| "collapsed" SurA (33.3) | o1s002 (17.3) | o1s016 (49.3) |
| "collapsed" SurA (47.7) | o1s003 (17.4) | o1s021 (34.8) |
| "collapsed" SurA (45.6) | o1s004 (21.1) | o1s016 (33.3) |
| "collapsed" SurA (32.2) | o1s006 (19.2) | o1s016 (48.6) |
| "collapsed" SurA (39.0) | o1s009 (34.1) | o1s016 (26.9) |
| "collapsed" SurA (46.7) | o1s009 (28.0) | o1s021 (25.4) |
| "collapsed" SurA (43.6) | o1s009 (52.1) | o2s012 (04.3) |
| "collapsed" SurA (33.3) | o1s010 (20.1) | o1s016 (46.6) |
| "collapsed" SurA (35.6) | o1s012 (30.8) | o1s016 (33.5) |
| "collapsed" SurA (26.2) | o1s013 (52.3) | o1s016 (21.5) |
| "collapsed" SurA (34.5) | o1s013 (43.7) | o1s021 (21.8) |
| "collapsed" SurA (43.4) | o1s013 (32.7) | o1s022 (23.9) |
| "collapsed" SurA (33.9) | o1s013 (58.8) | o2s006 (07.3) |
| "collapsed" SurA (25.6) | o1s013 (71.0) | o2s012 (02.7) |
| "collapsed" SurA (36.8) | o1s013 (61.7) | o4s001 (01.5) |
| "collapsed" SurA (34.5) | o1s015 (21.8) | o1s016 (43.7) |
| "collapsed" SurA (44.5) | o1s016 (43.1) | o1s017 (12.3) |
| "collapsed" SurA (46.7) | o1s016 (38.5) | o1s019 (14.9) |
| "collapsed" SurA (43.4) | o1s016 (43.3) | o1s020 (13.3) |
| "P2 closed" SurA (49.8) | o1s001 (08.7) | o1s016 (41.5) |
| "P2 closed" SurA (45.6) | o1s002 (10.5) | o1s016 (43.9) |
| "P2 closed" SurA (32.2) | o1s005 (16.5) | o1s016 (51.3) |
| "P2 closed" SurA (44.5) | o1s006 (11.4) | o1s016 (44.0) |
| "P2 closed" SurA (48.8) | o1s007 (14.8) | o1s016 (36.4) |
| "P2 closed" SurA (49.8) | o1s009 (19.9) | o1s016 (30.3) |
| "P2 closed" SurA (45.6) | o1s010 (12.3) | o1s016 (42.1) |
| "P2 closed" SurA (47.7) | o1s012 (18.3) | o1s016 (34.0) |
| "P2 closed" SurA (37.9) | o1s013 (36.9) | o1s016 (25.2) |
| "P2 closed" SurA (47.7) | o1s013 (28.7) | o1s021 (23.5) |
| "P2 closed" SurA (32.9) | o1s013 (63.8) | o2s012 (03.3) |
| "P2 closed" SurA (45.6) | o1s015 (14.0) | o1s016 (40.4) |

Each row of the table represents a combination of structures whose predicted scattering curves fit the experimental 0% SANS dataset (reduced chi-sq. < 1.05). Model 1 is always a conformation of SurA, which are detailed in Table S2. Models 2 and 3 are models of the SurA-uOmpA$_{171}$ complex. Numbers in parentheses represent the weight percentage of each model in the triplet used to fit the experimental SANS data.

***Table S8. Populations of Each Model in the SurA-uOmpA$_{171}$ Sparse Ensemble.***

| Model Name | Population (percent) |
|---|---|
| “P1 closed” SurA | 2.51 |
| “Collapsed” SurA | 23.04 |
| “P2 closed” SurA | 15.09 |
| o1s001 | 0.63 |
| o1s002 | 0.80 |
| o1s003 | 0.50 |
| o1s004 | 0.60 |
| o1s005 | 0.88 |
| o1s006 | 0.88 |
| o1s007 | 0.42 |
| o1s009 | 3.83 |
| o1s010 | 0.92 |
| o1s012 | 1.40 |
| o1s013 | 15.50 |
| o1s015 | 1.02 |
| o1s016 | 26.47 |
| o1s017 | 0.35 |
| o1s019 | 0.42 |
| o1s020 | 0.38 |
| o1s021 | 3.02 |
| o1s022 | 0.68 |
| o2s006 | 0..21 |
| o2s012 | 0.31 |
| o3s003 | 0.08 |
| o4s001 | 0.04 |

***Table S9. Contrast Values for Experimental Components.***

| % $D_2O$ | I(0) ($cm^{-1}$) | $C_{TOT}$ ($10^{-3}$ g $cm^{-3}$) | Contrast SurA ($10^{10}$ $cm^{-3}$) | Contrast 80% deut OmpA ($10^{10}$ $cm^{-3}$) |
|---|---|---|---|---|
| 0 | 0.22 | 3.0 | 2.4 | 5.8 |
| 30 | 0.054 | 3.0 | 0.8 | 4.1 |

### Supplemental Dataset Legends

***Dataset S1 (separate file) – paf_SurA_OmpA.xlsx***

This supplementary file contains all of the individual $uOmpA_{171}$ peptides that crosslinked to the eight high crosslinking efficiency $SurA_{pAF}$ variants, as determined by XL-MS. Each tab contains the results from each individual injection onto the Orbitrap, as well as a "Combined" tab, which contains a summary of all of the crosslinks found in the other tab.

***Dataset S2 (separate file) – paf_SurA_OmpX.xlsx***

This supplementary file contains all of the individual uOmpX peptides that crosslinked to the eight high crosslinking efficiency $SurA_{pAF}$ variants, as determined by XL-MS. Each tab contains the results from each individual injection onto the Orbitrap, as well as a "Combined" tab, which contains a summary of all of the crosslinks found in the other tab.

***Dataset S3 (separate file) – DSBU_SurA_OmpA.xlsx***

This supplementary file contains all of the crosslinks found between $uOmpA_{171}$ and WT SurA using the exogenous, chemical crosslinker DSBU. The "Total" tab contains the results from each individual injection onto the Orbitrap, and the "Combined" tab, which contains a summary of all of the crosslinks found in the other tab.

***Dataset S4 (separate file) – DSBU_SurA_OmpX.xlsx***

This supplementary file contains all of the crosslinks found between uOmpX and WT SurA using the exogenous, chemical crosslinker DSBU. The "Total" tab contains the results from each individual injection onto the Orbitrap, and the "Combined" tab, which contains a summary of all of the crosslinks found in the other tab.

***Dataset S5 (separate file) – DSBU_LCSurA_OmpA.xlsx***

This supplementary file contains all of the crosslinks found between $uOmpA_{171}$ and the "locked closed" (P61C/A218C) SurA variant using the exogenous, chemical crosslinker DSBU. The "Total" tab contains the results from each individual injection onto the Orbitrap, and the "Combined" tab, which contains a summary of all of the crosslinks found in the other tab.

***Dataset S6 (separate file) – SASD_scores_clustering.xlsx***

This supplementary file contains the solvent accessible surface distances (SASD) calculated for each pair of DSBU crosslinked residues found in the WT SurA-uOmpA$_{171}$ DSBU crosslinking experiments, mapped onto each of the 40 SurA-uOmpA$_{171}$ structural models. These distances were calculated using the Jwalk server and report on the ability of a structural model to accommodate an experimentally determined crosslink (small SASD means greater chance that a crosslink would form in this particular SurA-uOmpA$_{171}$ conformation (20). SASDs were used as inputs for a spectral biclustering analysis, allowing for different clusters of structural models representing distinct binding modes to be determined ("scores" tab).

**Supplemental Information References**

1. S. C. Gill, P. H. von Hippel, Calculation of protein extinction coefficients from amino acid sequence data. *Anal. Biochem.* **182**, 319–326 (1989).
2. N. R. Zaccai, *et al.*, "Deuterium Labeling Together with Contrast Variation Small-Angle Neutron Scattering Suggests How Skp Captures and Releases Unfolded Outer Membrane Proteins" in *Methods in Enzymology*, (Academic Press Inc., 2016), pp. 159–210.
3. K. L. Sarachan, J. E. Curtis, S. Krueger, Small-angle scattering contrast calculator for protein and nucleic acid complexes in solution. *J. Appl. Crystallogr.* **46**, 1889–1893 (2013).
4. J. E. Curtis, S. Raghunandan, H. Nanda, S. Krueger, SASSIE: A program to study intrinsically disordered biological molecules and macromolecular ensembles using experimental scattering restraints. *Comput. Phys. Commun.* **183**, 382–389 (2012).
5. A. V. Semenyuk, D. I. Svergun, GNOM - A program package for small-angle scattering data processing. *J. Appl. Crystallogr.* **24**, 537–540 (1991).
6. J. Trewhella, *et al.*, 2017 publication guidelines for structural modelling of small-angle scattering data from biomolecules in solution: An update. *Acta Crystallogr. Sect. D Struct. Biol.* (2017) https:/doi.org/10.1107/S2059798317011597.
7. SASSIE-Web, SASSIE-Web. *https://sassie-web.chem.utk.edu/sassie2/*.
8. M. C. Chambers, *et al.*, A cross-platform toolkit for mass spectrometry and proteomics. *Nat. Biotechnol.* (2012) https:/doi.org/10.1038/nbt.2377.
9. M. Götze, *et al.*, Automated assignment of MS/MS cleavable cross-links in protein 3d-structure analysis. *J. Am. Soc. Mass Spectrom.* **26**, 83–97 (2014).
10. J. M. A. Bullock, J. Schwab, K. Thalassinos, M. Topf, The importance of non-accessible crosslinks and solvent accessible surface distance in modeling proteins with restraints from crosslinking mass spectrometry. *Mol. Cell. Proteomics* **15**, 2491–2500 (2016).
11. A. N. Calabrese, *et al.*, Inter-domain dynamics in the chaperone SurA and multi-site

binding to its outer membrane protein clients. *Nat. Commun.* **11**, 1–16 (2020).
